# Scvi-hub: an actionable repository for model-driven single cell analysis

**DOI:** 10.1101/2024.03.01.582887

**Authors:** Can Ergen, Valeh Valiollah Pour Amiri, Martin Kim, Aaron Streets, Adam Gayoso, Nir Yosef

## Abstract

The accumulation of single-cell omics datasets in the public domain has opened new opportunities for reusing and leveraging the vast amount of information they contain. Such uses, however, are complicated by the need for complex and resource-consuming procedures for data transfer, normalization and integration that must be addressed prior to any analysis. Here we present scvi-hub: a platform for efficiently sharing and accessing single-cell omics datasets using pre-trained probabilistic models. We demonstrate that scvi-hub allows immediate access to a slew of fundamental tasks like visualization, imputation, annotation, outlier detection, and deconvolution of new (query) datasets, using state of the art algorithms and with a requirement for storage and compute resources that is much lower compared to standard approaches. We also show that the pre-trained models enable efficient analysis and new discoveries with existing references, including large atlases such as the CZ CELLxGENE Discover Census. Scvi-hub is built within the scvi-tools open source environment and integrated into scverse. It provides powerful and readily available tools for utilizing a large collection of already-loaded datasets while also enabling easy inclusion of new datasets, thus putting the power of atlas-level analysis at the fingertips of a broad community of users.

## INTRODUCTION

Machine learning models have been central to efforts to catalog cell states in health and disease with single-cell omics technologies (Kharchenko 2021; Wagner, Regev, and Yosef 2016; Heumos et al. 2023). These models are able to perform a variety of analysis tasks including dimensionality reduction, differential expression comparison, automated cell type annotation, denoising, deconvolution of spatial data, and modality imputation (Heumos et al. 2023). With the growth of single-cell data corpora, transfer learning will be an essential technique for accomplishing such tasks by leveraging large-scale datasets as reference atlases in a computationally efficient and performant manner. At present, transfer learning is primarily used for projecting cells onto a common low-dimensional representation that is employed for tasks such as annotation or trajectory inference (Lotfollahi et al. 2022; Kang et al. 2021; Hao et al. 2022). However, moving forward, additional applications will become more prevalent, such as interpretation of spatial data (Lopez et al. 2022) prediction of perturbation outcome (Roohani, Huang, and Leskovec 2023), prediction of multi-modal information from single modality data (Ashuach et al. 2023), detection of anomalous cellular subsets (Dann et al. 2023), or more robust analysis of differential expression (Boyeau et al. 2023).

Methods for transfer learning in single-cell omics broadly fall into two categories: non-parametric and parametric. In the non-parametric case, the algorithm uses the reference data directly to remove unwanted sources of variation. For example, Seurat and FastMNN integration utilize mutual nearest neighbors between reference and query data to remove query-specific effects (Hao et al. 2022; Haghverdi et al. 2018). In the second case, the algorithm uses a parametrized model like a conditional variational autoencoder (cVAE) to reduce the dimension of the reference data as well as remove unwanted variation. This same model can then be leveraged in order to project new data onto the same low-dimensional space. This approach is used by methods like scVI+scArches (Lotfollahi et al. 2022; Lopez et al. 2018; Gayoso et al. 2022), and can be efficiently designed, so that at query time only the query dataset is used, and not the original, potentially large, reference dataset. In many cases, these models also offer the benefit of a neural network decoder that can regenerate normalized raw count data with high fidelity from the low-dimensional representation. Thus, in principle, they can be used to analyze the reference data in its original dimension (e.g., at the gene level), when only provided with a much smaller (lower dimension) representation of that data

While parametric approaches have successfully been used in large-scale analyses (Suo et al., n.d.; Jones et al. 2022), there remain challenges with realizing the power of reuse of trained models. First, models can be trained using a variety of machine learning libraries and frameworks or use different versions of the same library, making them difficult to quickly ingest and use. Second, there is no standard for how pre-trained models should be deposited, which limits reuse to knowledge of a particular publication or model deposition. Finally, it can be difficult to assess the quality of a pre-trained model without standardized summary statistics of key performance metrics.

To address these issues, we introduce scvi-hub, a platform for sharing and re-using single-cell machine learning models that are implemented in the scvi-tools codebase (Gayoso et al. 2022). As a new component of scvi-tools, scvi-hub provides access to various popular single-cell model architectures spanning the core data analysis tasks, as well as to any future models that will be implemented using the developer API of scvi-tools (e.g., (Weinberger, Lin, and Lee 2023; Kleshchevnikov et al. 2022; Lotfollahi et al. 2022)). Scvi-hub facilitates model sharing through the Hugging Face Model Hub which offers an efficient, version-controlled, and feature-rich programmatic and user-facing interface. We developed scvi-hub with privacy preservation in mind and scvi-hub works with models deposited in cloud-based buckets like AWS S3. Scvi-hub provides functionalities that cater to both model consumers and model contributors. To the former community, it offers streamlined access to a variety of downstream analysis tasks using the downloaded models. Many of the downstream tasks are also made available in a form that relies on a low-dimensional (minified) version of the reference data (rather than the complete count matrix). This leads to substantial decrease in requirements for data storage and download bandwidth, thus increasing accessibility and inclusion within the data analysis community. Scvi-hub also offers a collection of tools to enable comparison and evaluation of models prior to uploading (by contributors) or prior to use of models for a specific analysis (by consumers) as well as a streamlined API for model upload and update (for contributors).

We demonstrate the utility of scvi-hub in a range of applications. One set of use cases considers reference-based analysis of query datasets - for annotation, outlier detection, and deconvolution. A second set of use cases considers only the reference, demonstrating the merit of tissue atlases as resources for making new discoveries. We have seeded scvi-hub with a collection of more than 90 models pre-trained on a variety of tissues and experimental conditions from the Tabula Sapiens consortium (Jones et al. 2022) and other large projects (see https://huggingface.co/scvi-tools). Scvi-hub also provides fully-featured access to the CELLxGENE census collection-the largest corpora of single cell omics datasets assembled to date (mouse and human data at s3://cellxgene-contrib-public/models/scvi). Scvi-hub is available as open-source under scvi-tools.org.

## RESULTS

### Scvi-hub facilitates reuse of machine learning models pre-trained on single-cell datasets

For the community of contributors, scvi-hub provides features that facilitate both evaluation and sharing of models (**Fig. 1**). Since all core models in scvi-tools are generative (i.e., capable of simulating new data), scvi-hub uses posterior predictive checks (Gelman, Meng, and Stern 1996) with model-simulated data to evaluate how well it captures the characteristics of the original input data. Here we implemented previously described metrics for single-cell omics data, including the coefficient of variation (Levitin et al. 2019; Gayoso et al. 2021), as well as new differential-expression-based metrics (see Methods). Importantly, these metrics are dataset-agnostic, in that they do not require dataset-specific information (such as cell-type labels or sample metadata) and are therefore broadly and immediately applicable.

**Figure 1:**
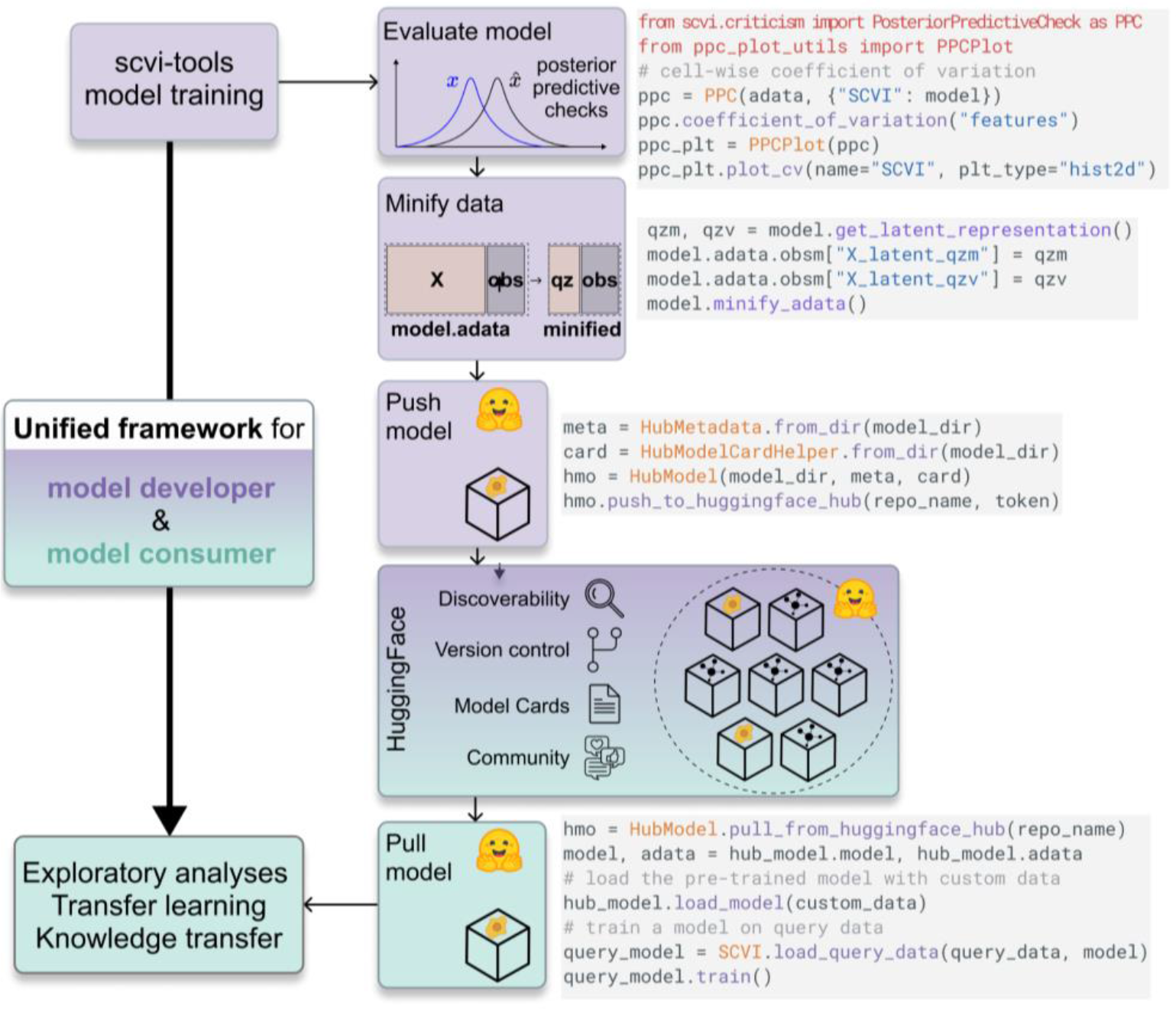
Overview of scvi-hub, depicting the new functionality for model contribution (purple) and retrieval (teal) that is implemented in scvi-hub, and its placement in an analysis workflow. A code snippet is presented for each of the tasks, highlighting the simplicity of this workflow. The workflow for model deposition (for model contributors) consists of training the model on reference data, evaluating model fit, minifying the reference data, and uploading the model to the scvi-hub. The training data (minified or raw) can be uploaded as well, either to Hugging Face or using any other online storage system (e.g., Zenodo) that can provide a direct link to access the data. This link is included in the Model card (visible through Hugging Face). The workflow for model retrieval (for the model consumer) consists of selecting the desired reference on scvi-hub and using the scvi-hub API for download and analysis. Note that the model consumers do not need to train a model on the reference data, and can also use the minified reference data instead of the raw form, thus lowering the current resource barriers (compute power, time and memory) for analysis. A single line of code switches from the minified version of the data to the full uploaded raw data to allow further downstream analysis.

To share models, scvi-hub leverages the Hugging Face Model Hub, which is a popular platform for sharing and using machine learning models from diverse domains, including natural languages, images, and audio. The Hugging Face Model Hub has features that make it ideal for single-cell genomics. One component of this is model discoverability, enabled via an advanced search interface and a uniform documentation and presentation with Model cards (description files that accompany and provide information on the uploaded model). It also comes with social features such as a discussion forum, which helps facilitate communication between contributors and consumers. Finally, the Hugging Face Model Hub provides features for maintaining consistency and backward compatibility through built-in git-based version control.

Model contributors also have the option to upload and share the data behind their model, thus allowing for a wide variety of uses (examples in **Fig. 2-3**). This can be done directly through Hugging Face or, alternatively, through any online storage system that can provide a direct link to access the data (e.g., Zenodo or CELLxGENE Discover). The data can be uploaded to Hugging Face either in its raw form (count matrix) or in a substantially reduced form, using a new feature we refer to as “data minification”. With this feature one can upload only the posterior parameters of the data, which are normally of a dimension much lower than the original count data. These posteriors can then be converted into an approximated and normalized form of the original data, using the generative part of the model. Since the models themselves often occupy orders of magnitude less space than the raw data, this feature makes available a compressed representation of a reference dataset, while still enabling much of the same functionality as with the raw data (**Supplementary Fig. 1**). As an example, in scVI, which is based on a cVAE, the minified data would be the latent space coordinates (mean and variance) of each cell. This representation (which is normally of less than 30 dimensions) can be converted into an approximated, and normalized form of the original gene expression data (genes used for model training, which are normally thousands) using the decoder part of the cVAE and enable efficient downstream analysis such as differential expression, feature correlation, and missing data imputation (Lopez et al. 2018; Gayoso et al. 2021; Ashuach et al. 2023; Boyeau et al. 2023; Steier et al. 2023).

**Figure 2:**
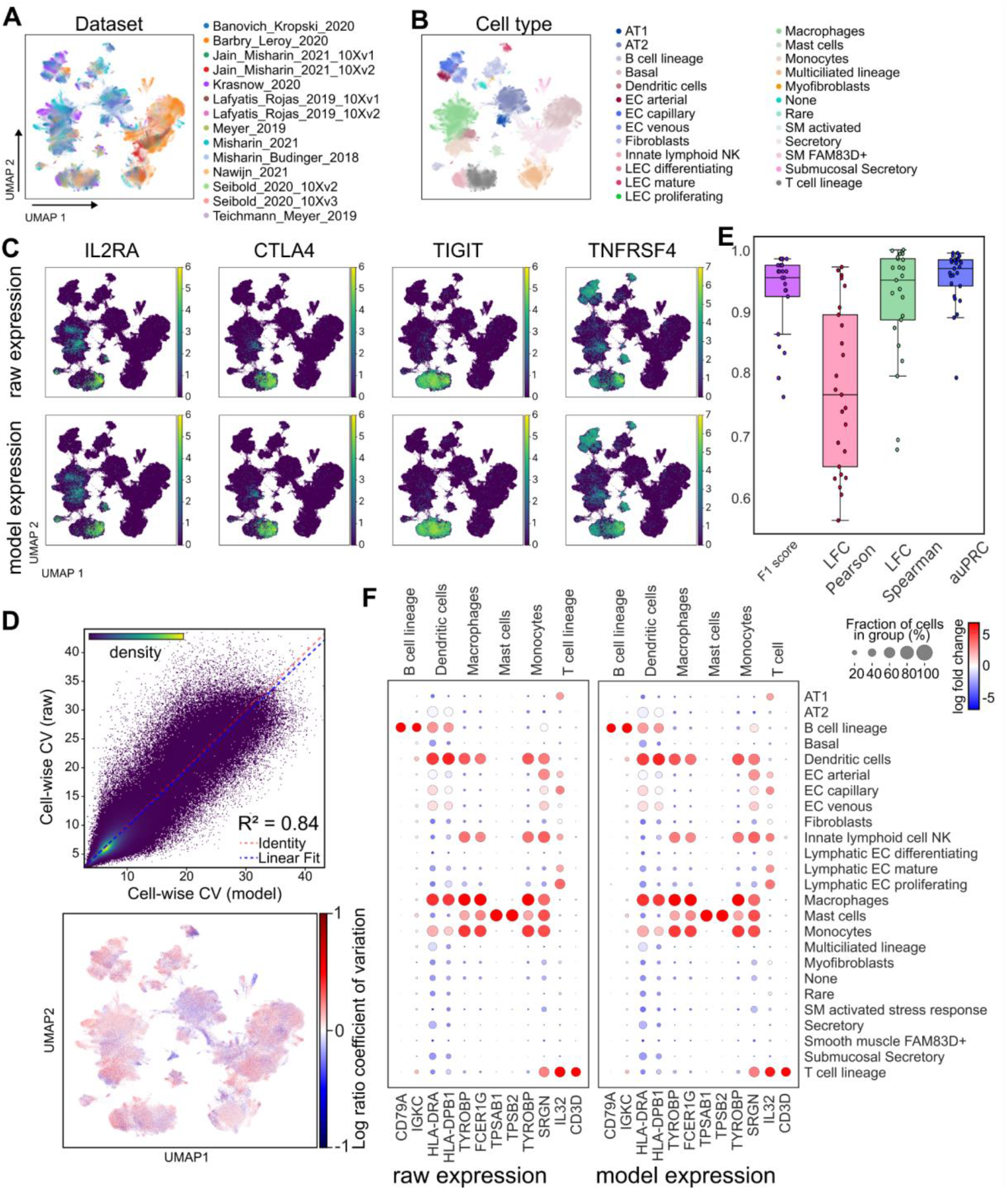
Reference-only tasks enabled by scvi-hub. Using the Human Lung Cell Atlas (HLCA) (Sikkema et al. 2022) as an example for a reference dataset in scvi-hub. The scANVI model object and a minified (latent space only) representation of the HLCA were used for common downstream tasks. **(A-B).** Visualization with UMAP, with cells colored by meta-data labels of interest (here - data source indicating the original studies included in the HLCA, and cell type). **(C).** Gene expression data can be regenerated from the minified data through the decoder network of the scANVI model. A single posterior predictive sample is used to estimate model expression. Expression levels of canonical regulatory CD4+ T cell markers are displayed after library-size normalization and log1p-transformation. Regulatory CD4+ T cells are not annotated in the HLCA and represent a fine-grained cell-type. **(D).** Coefficient of variation (CV) of the HLCA scANVI model. (Top) Each dot is a cell. The y axis reports the cell-wise CV computed on the raw data. The x axis reports the same metric computed on the generated data. (bottom) Discrepancy (Log-2 ratio) between cell-wise CV evaluated on model-generated counts versus raw data, overlaid on the UMAP embedding (Method). **(E).** Posterior predictive checks using differential expression between the cell-types in panel B (one vs. all); each dot represents metrics for one cell-type. Summary metrics from running Scanpy’s one-vs-all differential expression on the scANVI model-generated counts compared to the same metric computed on the original count data. Reported are: F1 accuracy, evaluated by overlap of the top 100 genes (using raw data vs. generated data for differential expression analysis), Pearson and Spearman correlation coefficients between the log-2 fold-changes (LFC) evaluated with raw vs. generated counts, and area under the precision-recall curve (auPRC) with genes identified by analysis of the raw data (adjusted p-value below 0.2) as the set of true hits and gene ranking defined by p-values evaluated with the generated counts. **(F).** Comparison of top marker genes for all immune cell-types (taking the top two for each type). Entries are colored by LFC of the respective one-vs-all comparison and sized by the number of cells with non-zero values for the respective gene in the raw data (left) or model-estimated proportion of expressing cells (right; see methods).

**Figure 3:**
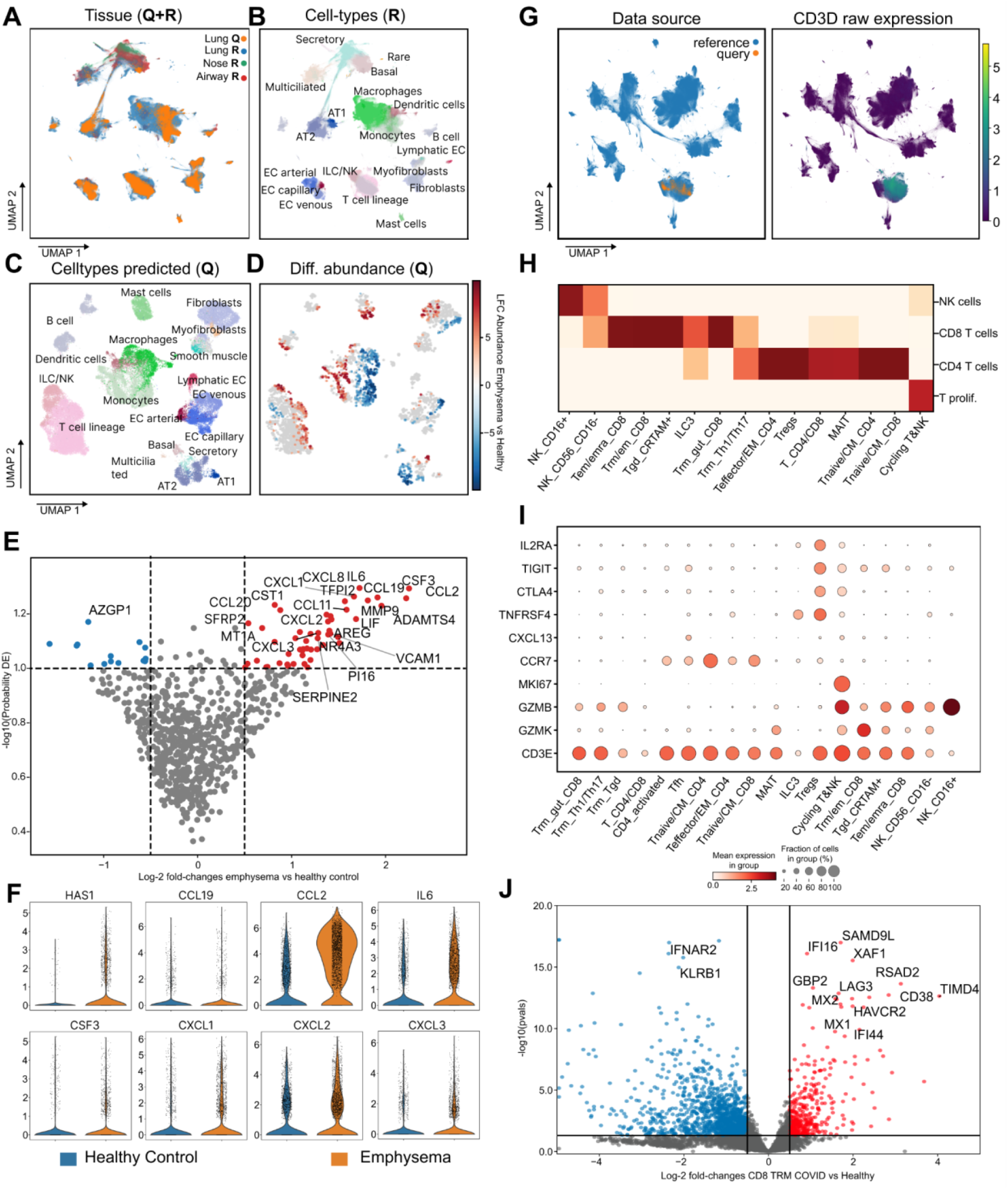
Query-to-Reference mapping tasks enabled by scvi-hub. **(A).** Joint embedding and UMAP visualization of the emphysema dataset (as query) and HLCA (as reference); cells are colored by their respective tissue and dataset. **Q** denotes the query dataset, while **R** denotes the reference dataset. **(B).** Visualization of the reference cells in the same UMAP coordinates, colored by coarse-level type (*ann_level_3* in HLCA). For display purposes, different annotations for smooth muscle cells and lymphatic endothelial cells were summarized to one label. **(C).** UMAP computed on query cells only (using their coordinates in the joint query and reference latent space). Cells are colored by their transferred cell-type (using summary of fine-grained labels to the annotation level of **(B)**; similar plot with fine-grained labels is presented in **Supplementary Fig. 5B**). **(D).** Differential abundance computed using Milo between cells from healthy donors vs. patients with emphysema in the query dataset. Plot is colored by the predicted log-2 fold-change for all neighborhoods with a significant change (FDR < 10%). **(E).** Model-based differential gene expression, comparing fibroblasts from the healthy vs. diseased samples in the query dataset. Mean log-2 fold-change is displayed on the x-axis and probability for non-DE on the y-axis. Top 20 genes based on their estimated significance are listed. All displayed genes are significant with an FDR < 10%. Genes with a model estimated mean <1e-4 in both groups are removed prior to display. **(F).** Violin plots of key differentially expressed genes based on the raw query data (library-size normalized and log1p transformed). **(G).** Embedding of T cells from the cross-tissue immune cell dataset (as query) integrated with HLCA reference dataset. Query cells almost entirely line up with CD3D+ reference cells (i.e., lymphoid cells, mostly T cells). CD3D expression is library-size normalized and log1p transformed. **(H).** Confusion matrix between the finest annotation scheme in HLCA (rows) and the labels infused using the query annotations (columns). **(I).** Hand-selected canonical marker genes for cell-types in panel H are displayed. Dotplots are computed based on the raw HLCA data (library-size normalized and log1p transformed). **(J).** Pseudo-bulk differential expression analysis between Tem/Temra_CD8 cells from COVID-19 infected samples vs. healthy controls using PyDESeq2.

Notably, the capability of working directly with minified data is becoming essential, as current atlas-level projects contain information on millions of cells, and analysis of those datasets has therefore been inherently tied to high performance computing resources. Data minification results in significantly less space requirements in memory (**Supplementary Fig. 1**), thereby enabling investigation of large-scale atlases on conventional hardware. It also increases computational speed since the latent embedding of the reference data (which is needed for any analysis that requires dimensionality reduction) is provided up-front and does not need to be computed anew.

The second set of users scvi-hub caters to are model consumers, who wish to analyze existing (reference) datasets or leverage reference datasets in order to analyze their own (query) data. Model consumers begin by searching and downloading their desired reference model and, if needed, its corresponding data in either minified or raw form. These steps can be done either through the Hugging Face website or directly through the scvi-hub Python API (which internally uses the Hugging Face API). Since scvi-hub is part of the scvi-tools package and the scverse ecosystem ((Virshup et al. 2023), which includes Scanpy (Wolf, Angerer, and Theis 2018)), consumers can seamlessly integrate the downloaded models into existing analysis workflows, which provide a slew of functionalities for exploration, visualization and analysis. Extensive tutorials exist to work with scvi-hub in the R ecosystem as well, thus supporting downstream analysis with Seurat (Butler et al. 2018) and other popular environments.

In the following sections, we describe the functionalities for model criticism, which can be employed by both contributors (before uploading their models) and consumers (to check existing models before conducting analysis and to check whether query data fits to a reference model). We then describe the functionalities that are available for consumers through their downloaded models, including both analysis of the reference data (using its minified form), and analysis of user-provided data. For the former, we also demonstrate how one can leverage knowledge from the user-provided data to refine the analysis of the reference dataset.

### Scvi-hub provides evaluation procedures to select models and test their suitability for query datasets

Evaluation of model quality is an essential part of scvi-hub. It allows contributors to scrutinize their models before upload, and consumers to verify that the models that they download are relevant and of sufficient quality. To achieve this, we developed *scvi.criticism* - a new module for evaluating models that were trained with scvi-tools. *scvi.criticism* implements posterior predictive checks (PPCs), which compare the distribution generated by the fitted model and the actual observed data (Gayoso et al. 2021; Lopez et al. 2018; Gelman, Meng, and Stern 1996). To conduct PPC, we first sample from the distribution of the data as predicted by the fitted model and then compute summary statistics per gene (coefficient of variation and differential expression between pre-specified groups of cells) on this distribution, as well as on the raw data. The PPC consists of measuring the closeness between the resulting pairs of statistics (raw vs. generated). Close similarity is a standard measure that defines a well-trained model that is representative of the original data. To demonstrate this, we computed PPC on the Human Lung Cell Atlas dataset (HLCA, (Sikkema et al. 2022)) using the author-provided scANVI model (Xu et al. 2021) that was trained on this data (**Fig. 2A-B**). We reported the coefficient of variation as well as a set of metrics computed based on the results of differential expression between the author provided cell-type labels, performed on the predicted data and on the raw data (**Fig. 2D-F and Supplementary Fig. 2**). We observed that the data generated by the model fits well to the raw data, e.g., identifying similar sets of differentially expressed genes in a one vs. all cell type comparison. We additionally used the model to generate gene counts from the minified data and found agreement between model-predicted expression and observed gene expression for multiple markers of regulatory T cells, which were not annotated as a separate cell-type in the HLCA. This shows that we can recover fine-grained cell types (potentially not annotated in the original reference data) by generating their corresponding marker expression using the minified data and the pre-trained model (**Fig. 2C)**.

Model contributors can use *scvi.criticism* to evaluate the goodness of fit of their models, select among several candidate models (e.g., with different hyperparameters), and optionally include these evaluations as part of the respective Model Card on Hugging Face. This is particularly useful in the case where the data is minified as it provides more confidence in the reliability of the counts generated by the model in the absence of the full raw counts. To demonstrate this, we used *scvi.criticism* to evaluate the goodness of fit of one well-trained and two poorly-trained scVI models on the Heart Cell Atlas dataset (Litviňuková et al. 2020) (**Supplementary Fig. 3** and Methods). We compared the cell-wise coefficient of variation results computed on the raw data and on the estimated data from each of the three models (**Supplementary Fig. 3A**). We observe that the well-trained model performs better (higher Pearson correlation with the raw coefficients of variation) than both of the poorly-trained models. We also report the similarities between the results of differential expression computed on the raw and estimated data from each of the three models (**Supplementary Fig. 3B**). We used multiple metrics for this comparison (Methods) and observed, once again, that the well-trained model performed consistently better than either of the two poorly-trained models in most metrics.

Another important use case of *scvi.criticism* is to evaluate the extent to which a reference model is apt for analyzing a query dataset (**Supplementary Fig. 4**). To demonstrate this, we created a query dataset consisting of all epithelial cells from the Tabula Sapiens project (spanning different tissues) and used the HLCA pre-trained scANVI model as our reference (i.e. many cell types, but airway-only). We projected the query dataset onto the reference model, using the scArches functionality (available on scvi-tools (Lotfollahi et al. 2022; Lopez et al. 2018; Gayoso et al. 2022)). We then fed the obtained latent representation of the query data to the generative part of the model after transfer learning to test whether it is capable of generating gene expression profiles that are similar to the raw query data (evaluated using *scvi.criticism*). Reassuringly, we see very good performance on lung epithelial cells, airway epithelial cells as well as epithelial cells from the back part of the tongue (arguably similar to upper airway epithelium). These organs are all included in the HLCA reference dataset. For tissues that are not represented in the HLCA, we find substantially worse correlation, indicating that the model does not capture their underlying distribution of gene expression well. Criticism therefore allows detecting reference models that are well suited for the query dataset at hand and rely on those for downstream analysis.

### Scvi-hub enables efficient exploration and re-analysis of large reference datasets

The first set of analyses facilitated by scvi-hub is centered on the reference datasets, without inclusion of any additional (query) information. This approach allows for rapid exploration and re-analysis of reference atlases (without requiring time-consuming and resource-heavy model training procedures), as well as efficient generation of new models that are based on these references. After searching for a model that fits the user’s specification on Hugging Face (i.e., tissue, cell types, and a modeling scheme, such as scVI, DestVI etc.), model consumers can pull (download) the model and its corresponding data, which can be either in raw or minified format. Analysis of reference datasets includes operations that concern the low-dimensional (latent) representation of cells, such as visualization, clustering, trajectory inference and differential abundance analysis. Additionally, model consumers can perform analyses at the high-dimensional, omics measurement level by accessing the raw data or by generating count data using the minified format. These high dimensional representations can then be readily analyzed in Scanpy (Wolf, Angerer, and Theis 2018) or Seurat (Butler et al. 2018), through a slew of procedures. As an example, using the HLCA reference scANVI model (which had good PPC performance; **Fig. 2D-F**), we find that re-analysis of model-generated gene expression profiles (using the minified data) can add much detail to the available annotations. For instance, we observe that regulatory T helper cells (Tregs), which is not a label included in the reference annotation, can be well identified based on generated expression levels of a few marker genes (*IL2RA*, *CTLA4*, *TIGIT*, *TNFRSF4*; **Fig. 2C**). We further probe this point in our analyses in **Fig. 3G-J**, described below.

Downloading minified data and the pre-trained model therefore allows preservation of information (compared to downloading raw data) but allows to rely on a much smaller dataset. This allows fast interaction with the data using all downstream capabilities offered by Scanpy or Seurat, and also all built-in functions in scvi-tools (e.g. for differential expression (Boyeau et al. 2023)). As scvi-hub stores the link to the full data object by invoking a single-line function call, the full data can also be downloaded and particular findings made on minified data can be validated on the full raw data.

### Scvi-hub enables efficient reference-based analysis of query data by transfer learning

The most fundamental part in analyzing a query dataset given the appropriate reference is to represent the query data using the reference model. In the context of scvi-tools, this is done by calculating the coordinates of each query cell in the latent space of cell states that is encoded by the model and that is initially populated by the reference cells (**Fig. 2B**). While the encoder networks in scvi-tools can readily provide latent representation for any query cells, these representations might be distorted by batch effects. An efficient way to address this (while not retraining the model anew on the reference and query data together – a resource-consuming procedure) is to perform minimal training only to capture these batch effects and leaving the model otherwise unchanged. This procedure, which is implemented by scArches (Lotfollahi et al. 2022), effectively produces a joint embedding of the reference and query dataset while quickly removing unwanted variation. Such joint representations can be leveraged for a variety of downstream tasks that are readily available through scvi-hub. Here, we demonstrate the capabilities of scvi-hub to analyze query data with transfer learning, through four such fundamental tasks: visualization, annotation, anomaly detection, comparative analysis, and deconvolution.

As a first test case, we used a query dataset of three healthy individuals and three emphysema patients (Wang et al. 2023) and designated the HLCA as reference. The two datasets (query and reference) are well-integrated with scANVI as the reference model and with use of scArches to add in the query data, leading to an informative, reference-informed, visualization of the query data (**Fig. 3A-B**). To annotate the states of cells in our query, we transferred labels from HLCA using a simple KNN classifier (using the reference embedding in the joint latent space to identify K-neighborhoods; **Fig. 3b** and **Supplementary Fig. 5**). We found that the reference-based annotation is consistent with the labels provided by the original study of the query dataset, yet adds a great deal of resolution. For instance, cells that were originally labeled as Endothelial cells (EC) were now divided into several sub-groups of that lineage, including venous systemic, venous capillary, arterial and aerocyte capillary (**Supplementary Fig. 5A**-B).

The reference-based probabilistic representation computed for each query cell can also facilitate comparative analysis within the query dataset. To explore this, we first used Milo (which relies on the integrated embedding space) to compare the composition of cell states in the three healthy query samples versus the three emphysema-affected samples. We found a significant (FDR < 10%) increase in the abundance of certain states of macrophages, fibroblasts and epithelial cells in the emphysema samples (**Fig. 3D** and **Supplementary Fig. 6A**-B). To get more insights on the shift in cell states, we next used the differential expression function built in scvi-tools (which uses the reference-based scVI model) to explore disease-associated gene expression changes in fibroblasts of the query data (**Fig. 3E**). We find that fibroblasts in emphysema patients strongly up-regulate pro-inflammatory chemokines that attract neutrophils (*CXCL1/CXCL2/CXCL8*), monocytes (*CCL2, CSF3*) and naive T cells (*CCL19, CCL20*).

The original publication highlighted the role of fibroblasts in inducing a niche for resident memory T cells, further corroborated by the fact that depletion of fibroblasts in a mouse model led to decrease of resident Th17 cells. Our reference-powered analysis therefore highlights additional putative mechanisms associated with neutrophils and monocytes. Indeed, neutrophils release granule proteins like neutrophil elastase and myeloperoxidase and have thus been associated with emphysema (Gernez, Tirouvanziam, and Chanez 2010). Similarly, monocytes-derived macrophages produce metalloproteinases leading to tissue remodeling in emphysema (Shapiro 2012). To evaluate the importance of using a reference-based model for achieving these insights (despite the low number of samples), we conducted a pseudo-bulk differential expression analysis, comparing emphysema vs. control fibroblasts using PyDESeq2 (Muzellec et al. 2023) as well as using scvi-tools differential expression function after training a scANVI model trained from scratch but initializing the weights with the reference-based model. In both cases we were not able to identify the same clear signatures that we identified using scvi-tools leveraging the reference model (Supplementary Fig. 7, Supplementary Fig. 8). We additionally validated our findings by displaying the measured gene expression (from the query data only) of the genes that were identified by the reference-based analysis (**Fig. 3F** and **Supplementary Fig. 6C**).

Another area of single cell genomics where a form of transfer learning can be used to gain new insight is spatial transcriptomics (ST). Since the resolution of most sequencing-based ST protocols is at a level coarser than a single cell, deconvolution is needed to estimate the cellular composition of each measurement unit (often referred to as a spot). Algorithms such as DestVI (Lopez et al. 2022) and Stereoscope (Andersson et al. 2020) were developed to leverage models that are pre-trained on non-spatial scRNA-seq datasets to perform such deconvolutions. In contrast to most other algorithms, DestVI and Stereoscope (both available on scvi-hub) allow pre-training a first model on a reference scRNA-seq data and then, in a separate procedure, apply this model to deconvolve any incoming ST data. Scvi-hub users can thus download the scRNA-seq parts of DestVI and Stereoscope models that were pre-trained on the relevant atlas (i.e., of the same tissue) and deconvolve their spatial data. We demonstrate this using a spatial dataset of a human prostate, generated with 10x Visium technology (“10X Visium Prostate” n.d.). We deconvolved this data (provided at a resolution of 50um) using a DestVI model pre-trained on the prostate section of the Tabula Sapiens reference atlas (Jones et al. 2022). The deconvolution paints a picture of cellular localization in the prostate that is consistent with the literature, with clear demarcation of prostate glands and stromal regions. Specifically, in the glands, we detect luminal cells of the prostate epithelium that are surrounded by fibroblasts. In stromal regions we detect smooth muscle cells (**Supplementary Fig. 9)**. Beyond deconvolution, the use of DestVI estimates cell-type specific gene expression inside each relevant spot, thus opening the way for downstream analysis such as comparing how cells of the same type differ between tissue areas (Lopez et al. 2022).

### Scvi-hub enables efficient re-analysis of reference data sets by infusing novel insights from query data sets

The joint embedding of query and reference datasets can also be used for an additional, less prevalent procedure, which we term label infusion. While reference atlases tend to include large numbers of samples and cells, their levels of annotations can be limited in granularity, at least for some compartments. Conversely, other more narrowly focused datasets may provide higher resolution, e.g. thanks to more targeted capture of certain cellular subsets or by more nuanced procedures for cell-state labeling. By reannotating reference datasets based on labels in the query datasets, fine cellular subsets of interest can therefore be identified and their function in (query) disease contexts can be better studied leveraging the power of the reference dataset.

We demonstrate label infusion by refining the cell-type labeling of NK and T cells in the HLCA dataset (**Fig. 3G**). We used a recent study of immune cells across different organs (Domínguez Conde et al. 2022), subsetted this study to only NK and T cells across all different organs, and integrated the resulting “query” dataset with the HLCA reference, using its scANVI model. After integration with scArches, we confirm that the query cells are well integrated with NK and T cells of the reference dataset. The joint embedding then helped us transfer knowledge from the query to the reference (label infusion; **Fig. 3H**). For instance, this analysis helped confirm the presence of regulatory T cells in the HLCA, highlighting the same set of cells that we reannotated in our reference-only analysis, using generated expression of marker genes (**Fig. 2C, Supplementary Fig. 9)**. We further validated other infused cell-type labels by checking their canonical cell-type marker genes (**Fig. 3I**). For instance, we confirmed that the cells relabelled as T follicular helper cells (Tfh) are distinguished by the expression of their canonical marker *CXCL13*.

The infused labels can be used to gain new insight from the reference data. To demonstrate this, we focused on a subset of cells that we reannotated as CD8+ resident memory T cells (labeled as Tem/Temra_CD8 in **Fig. 3**). Since the HLCA includes samples from COVID patients in addition to healthy donors, we were now able to look at the specific effects of infection on this more narrowly defined immune subset. This analysis finds an upregulation of markers of exhaustion of CD8+ T cells (*LAG3, CD38, TIMD4*, and *HAVCR2*) as well as increase of Interferon-regulated genes (*GBP2, MX1, MX2, IFI16, XAF1, SAMD9L,* and *IFI44*), both of which are consistent with recent, more targeted, studies of COVID infection (Szabo et al. 2021; Rha and Shin 2021).

### Scvi-hub provides a scalable access and efficient analysis environment for very large references: the case of CZ CELLxGENE Discover Census

Recently, several large corpora of single cell omics datasets were collected to enable comprehensive view of cell types and states in various tissue contexts, thus starting to realize the potential of the cell atlases <u>(Regev et al. 2018)</u>. The largest of these efforts is the CELLxGENE census collection (CZI Single-Cell Biology Program et al. 2023). It currently contains above 30 million human cells from a broad array of tissues, diseases and experimental settings. Leveraging large collections such as the Census removes the necessity to find a suitable reference dataset. Instead, a query dataset can be compared against the whole existing census, in a manner reminiscent of searching DNA sequences in a reference genome. Furthermore, Census-level data collections provide an attractive resource for analysis by themselves, enabling one (to name a few examples) to identify and study new cell subsets, gene expression programs, and gene-gene dependencies.

To demonstrate how scvi-hub can help realize the potential of census-level analyzes, we leverage an scVI model that was trained to embed the census collection. This model is created and managed by the CELLxGENE Census team; it is being routinely updated to include new datasets,. This model is also accessible by scvi-hub and its full array of features, thus opening the way for users to leverage this ever-growing resource that spans diverse populations, tissues, disease contexts and cell subsets (https://chanzuckerberg.github.io/cellxgene-census/). A first clear advantage of a model-based representation of the Census collection is in the volume of the data. While the current release of the Census collection requires 500 GB of disk and hours of download time, the minified form of this data requires 30 GB of disk space and (in our hands) less than 30 minutes for download. This reduction the size allows fast exploration and interaction with the Census and significantly reduces the entry barrier (in terms of compute resources and internet bandwidth) for leveraging this potentially transformative resource.

As our case study, we used as query a dataset from a study of anti-CD19 CAR therapy (autologous axicabtagene ciloleucel; <u>(Deng et al. 2020)</u>), which was not included in the Census at the time of analysis. In this study, infusion bags from 24 treated patients were collected and single-cell RNA-seq was performed on the cell suspensions inside each bag. These suspensions consist primarily of CAR-T cells, spanning a heterogeneous set of states. Coupling information on treatment responses with knowledge of cell states provides a promising strategy for prognosis, improvement of the CAR products, and prediction of adverse effects [REF].

We used the scArches functionality that is built in scvi-hub to to efficiently integrate the query dataset with the Census collection. The use of scArches greatly simplified this process since learning the integrated model only required processing the query cells (and not the reference collection of 30M cells). Using the model criticism package of scvi-hub, we found an encouraging correlation between the coefficient of variation of the query cells, comparing the model-based values to the raw data (**Fig. 4A**). Within this analysis, we found two cell subsets with a lower agreement of the coefficient of variation and identified those as cells with a high fraction of mitochondrial reads and nuclear RNA, indicating low quality events (**Supplementary Fig. 11 A, B**). Looking at the integrated data we find that cells integrate well with immune cell subsets within the full census dataset. For annotation, we transferred labels from the Census to the query dataset using a KNN classifier defined on the integrated space. We restricted this analysis to only consider reference labels that originated from an organ-wide study of immune cells, recently incorporated into the Census (**Fig. 4B, C, Supplementary Fig. 11 C**) <u>(Domínguez Conde et al. 2022)</u>. We chose this dataset for label transfer thanks to its high level of annotation. Overall, we see that the transferred labels line up well in terms of the expression of marker genes (**Fig. 4D**). In addition, this reannotation adds granularity to the labels assigned by the original study of the reference dataset, while maintaining overall consistency (**Supplementary Fig. 11 D, E**). For instance, we observed a subset of cells, previously labeled as CD4+ T cells, that is now more finely labeled as regulatory T cells - an annotation that is supported by their expression of *FOXP3*. Similarly, a subset of cells previously assigned with the general term of CD8+ T cells are now specifically labeled as terminally differentiated (**Fig. 4D, Supplementary Fig. 11B)**.

**Figure 4:**
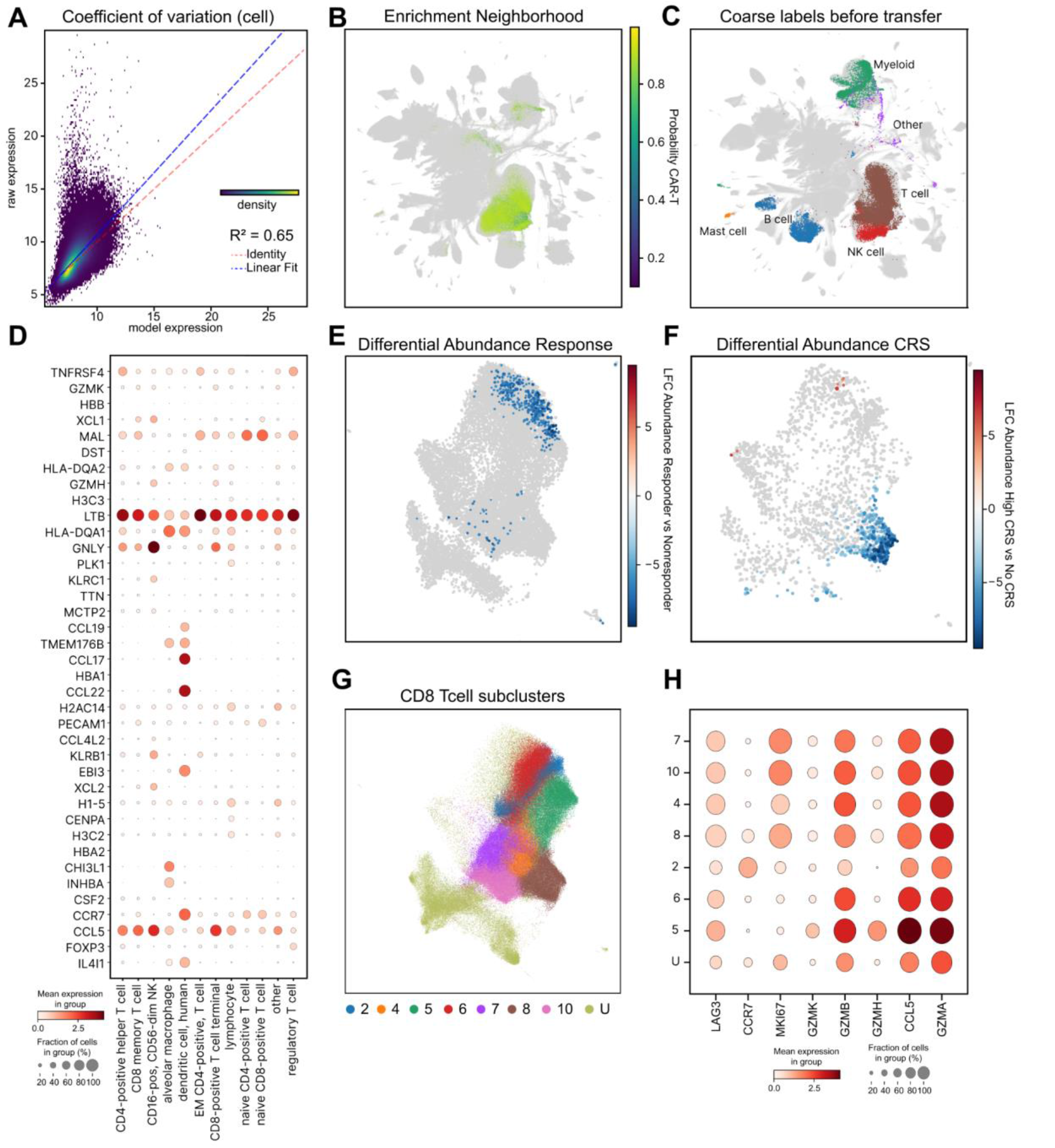
Query analysis using census-level pretrained model. Using a custom model trained on the CELLxGENE census data to analyze a dataset with CAR T cells. **(A)** Coefficient of variation on query data after transfer learning the pretrained model. Each dot is a cell. The y axis reports the cell-wise CV computed on the raw data. The x axis reports the same metric computed on the generated data. **(B)** Probability that the cell neighborhood contains CAR T cells displayed on a random subset of census data set. We concatenate the query data and the same amount of cells from the whole census data set. Cells are colored by the probability that cells in the neighborhood are query cells. Gray cells contained no cells from the query data set in the neighborhood graph. **(C)** Same embedding as in **b** colored by the cell-type labels of the cross tissue immune cell atlas <u>(Domínguez Conde et al. 2022)</u>. Coarse labels are written to annotate the colors. For a full legend view **Supplementary Fig. 11C**. Most CAR T cells overlap with regions with T cells and myeloid cells. **(D)** Marker genes computed on query cells. Marker genes were identified using differential expression function in scVI. Displayed is the percentage of cells expressing cells as well as the mean expression after library size normalization and log1p transformation of raw counts. **(E)** Milo differential abundance analysis overlaid on top of UMAP embedding of query cells computed on latent embedding of cells in the query model. Displayed is the estimated log-2 foldchange in abundance for all cell neighborhoods with an FDR < 10%. **(F)** Similar analysis, while estimating the effect size for patients without cytokine release syndrome (CRS) and with high grade (ICANS grade 3 and 4) CRS. Displayed is the estimated log-2 foldchange for cell neighborhoods with an FDR < 10%. **(G)** Leiden clustering of all clusters annotated manually as CD8 T cells. Cells in clusters with metrics of low quality are annotated as Other. **(H)** Hand selected marker genes are displayed to distinguish between different states of CD8 T cells. Cluster 8 is associated with no CRS and good response. Displayed is the percentage of cells expressing cells as well as the mean expression after library size normalization and log1p transformation of raw counts.

The census-embedded and re-annotated representation of the reference data opens a way to further study the association between responses to CAR therapy and the composition of the infusion products. To this end, we applied Milo <u>(Dann et al. 2022)</u> on the embedding coordinates of the query dataset to identify cell subsets that are associated with poorer responses. We found that terminally differentiated CD8+ T cells as well as regulatory T cells are associated with reduced response (FDR<10%; **Fig. 4E**). The findings on terminally differentiated CD8+ T cells align well with the results in the original study, which reported a subcluster C1 with markers of exhaustion in partial responders. The findings on Treg also accord with previous reports about negative association between their concentration in the circulating CAR population post-infusion, and the success of CAR therapy <u>(Good et al. 2022)</u>. Through our reanalysis of the query data enabled by integration with census data, we were able to track these cells back to the infusion product and establish their negative association with treatment outcome already at that early, pre-treatment stage. We additionally used milo to identify cell states that are significantly (FDR <10%) associated with cytokine release syndrome (CRS) which is the most common and potentially life threatening adverse reaction to CAR T therapy. We identified a subset of CD8+ T cells with high proliferation (*MKI67* positive) and expression of *CCR7* to be negatively associated with CRS (**Fig. 4F)**. In addition, these cells express higher levels of *GZMB*, *GZMA*, *LAG3* and *CCL5* compared to naive CD8+ T cells (Leiden cluster 2; **Fig. 4G-H** and **Supplementary Fig. 11D)** and correspond to early activated central-memory like CD8 T cells. We have therefore demonstrated that our reference-based analysis can uncover novel cell states associated with treatment response and negative side effects, while reducing manual optimization of various hyperparameters.

Surprisingly, our reannotation also identified a population of dendritic cells that are positive for *CCR7*, *CCL17* and *CCL22* as well as a subset of NK cells. Both of these subsets were not identified in the original study (**Fig. 4D** and **Supplementary Fig. 11E**). As far as the standard protocols go, dendritic cells were not meant to be included in the infusion products and it is unclear what relevance these cells have to CAR T treatment. However, overexpression of CCR4 on CAR T cells, which interacts with *CCL17* and *CCL22*, has been demonstrated to increase efficiency of T cell treatment <u>(Foeng et al. 2022)</u>, suggesting a potential beneficial effect in co-stimulation of T cells with activated donor-derived dendritic cells.

### Reannotating the CELLxGENE Discover Census enables fine-grained analysis of cell states in cancer

The CELLxGENE census contains datasets from different studies, which can have very different granularity of their cell-type annotations as well as different definitions (e.g., marker genes) for declaring some cell-types. Harmonizing those disparate annotations opens the way for studying cellular diversity in a large variety of physiological contexts and thus has the potential to enable insights at an unprecedented scale and breadth. Scvi-hub provides readily available tools to accomplish this, through its functionalities for reference data access and analysis. To demonstrate this, we used the labeling scheme of the cross-tissue immune atlas to re-annotate the remaining cells in the CELLxGENE census. To this end, we accessed the scVI model and the respective minified representation of the Census data and applied a KNN classifier to transfer labels of cells of the cross-tissue immune atlas to all other cells in the Census. We also used the KNN classifier to identify Census cells that could not be annotated with this approach (no labeled cells present in their K-neighborhood). This resulted in identification and reannotation of immune cells in the Census as well as exclusion of non-immune cells, as further supported by the expression of the common leukocyte marker *PTPRC* (**Fig. 5A, Supplementary Fig. 11C**).

**Figure 5:**
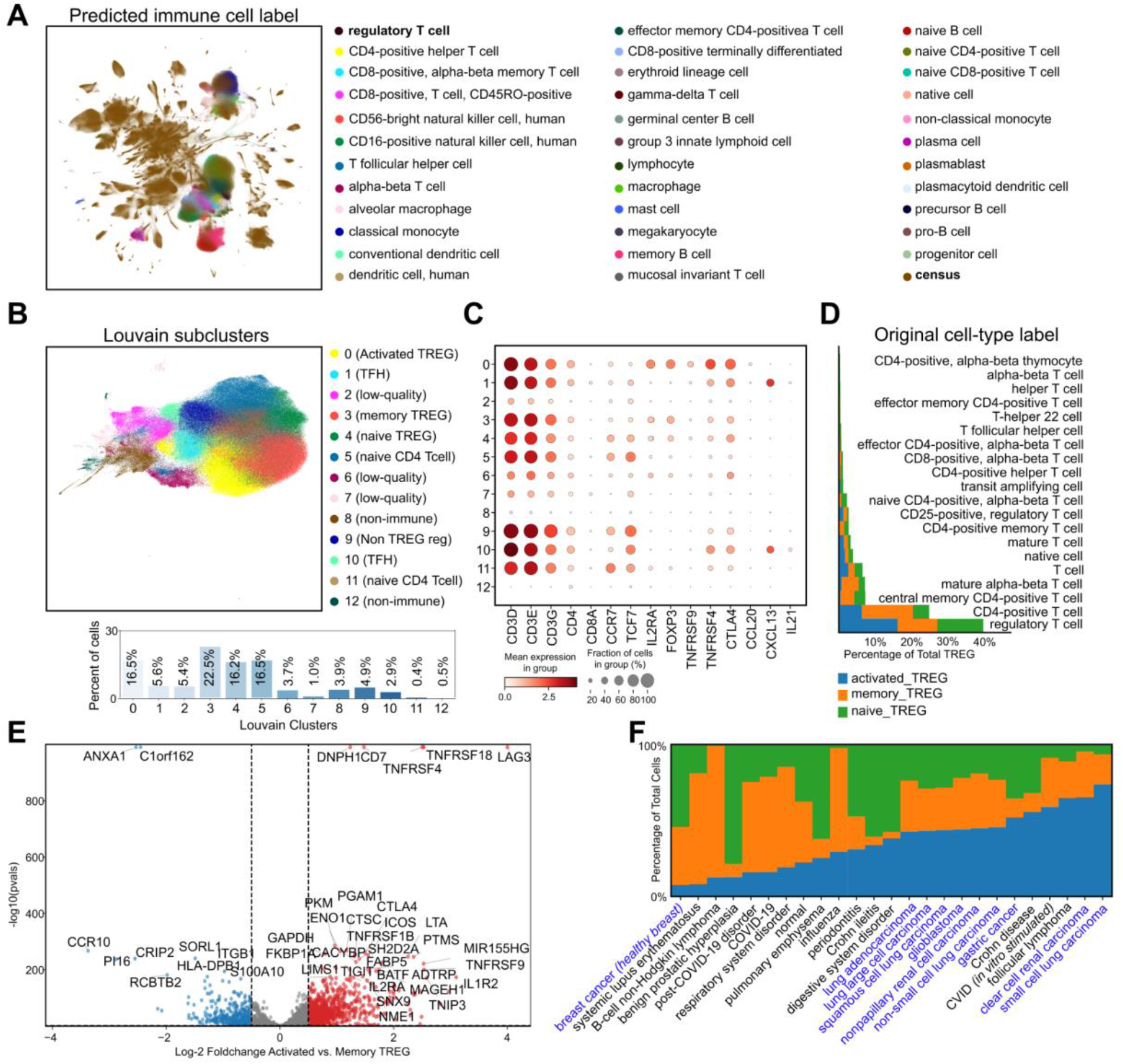
Refined analysis of CELLxGENE census using scvi-hub. We transfer labels from a well annotated immune cell dataset to all other datasets and reanalyze those. **(A)** Labels were transferred using a concatenation of cells from <u>(Domínguez Conde et al. 2022)</u> and the same amount of random census cells labeled “census”. Cells are annotated with an immune cell label if any cell in a neighborhood of 50 cells was annotated by this cell-type and is otherwise labeled “census”. Displayed is a random subset of all census cells as UMAP embedding of 36 million cells can’t be performed on our hardware. **(B)** Dataset was subset to all cells labeled TREG in **A** and another scVI model was trained using only these cells. Displayed is the UMAP embedding of these cells, Leiden clustering and our manual annotation of these cell states. (Bottom) Barplots of frequencies within each Louvain cluster in comparison to all cells annotated as regulatory T cell (cluster 0, 3, 4, 6 are regulatory T cells after manual verification and make up ∼60% of the annotated cells). **(C)** Hand-selected marker genes are displayed. Displayed is the percentage of cells expressing cells as well as the mean expression after library size normalization and log1p transformation of raw counts. **(D)** Cells were subset to Leiden cluster 0, 3 and 4 with marked expression of *FOXP3*. Displayed is the percentage of total cells annotated with a specific cell-type label in the CELLxGENE census data. Bars are colored based on the cell state based on Leiden clustering. **(E)** Pseudobulk differential expression analysis using PyDESeq2. As the sample ID we used the concatenation of donor_id, dataset_id, tissue and assay. Displayed is the comparison of cells annotated as Activated TREG vs cells annotated as memory TREG. Top-50 marker genes are displayed. **(F)** Displayed are all disease categories with more than 200 cells in Leiden clusters labeled TREG. We display the fraction of each TREG subtype for each disease. In blue we highlight all disease categories of solid tissue cancers.

As part of this analysis, we identified 370,000 cells as Tregs - a designation confirmed by relatively prevalent expression of *FOXP3* and *CD3D* (**Supplementary Fig. 12C**). For a more detailed analysis of these cells, we downloaded their raw expression profiles and retrained an scVI model. When downloading, we used the soma API provided by the Census project to access only the cells that we annotated as Tregs, thus substantially reducing the required time for download (30 minutes vs. 10 hours that are needed for the full Census dataset). Using the resulting embedding, we clustered the cells into groups and annotated them manually (**Fig. 5B-C**). Through this additional inspection, we found that approximately 40% of the cells likely represent other immune populations, such as follicular T helper cells that express regulatory molecules (e.g., *CTLA4*; Clusters 1 and 10; ) and naive CD4+ T cells (Clusters 5 and 11). The remaining 60% of cells were identified as Tregs that are organized into three major types (clusters 0, 3 and 4). The first type expressed high levels of *CCR7* and *TCF7* and thus labeled as ‘naive Treg’, the second type expressed high levels of *PI16* and *ANXA1* which we labeled as ‘memory Treg’, and the third type expressed high levels of *TNFRSF9*, *IL10* and *CTLA4* and labeled as ‘activated TREG’ (**Fig. 5B-C, Supplementary Fig. 13**). Interestingly, about 60% of these cells were not originally labeled (in their respective study) as Tregs, but rather with less specific labels such as ‘T cell’ or ‘CD4+ T cell’ (**Fig. 5D**).

This re-identification and re-labeling of Tregs in the Census collection opens the way for studying which states in this compartment are associated with specific tissues or diseases. To explore this, we first used the sample meta-data fields in the Census collection to designate each cell as coming from a healthy individual or from a study participant that is impacted by a certain pathology. Considering tissues that were profiled in healthy individuals, we found the highest proportion of activated Tregs (out of the entire Treg compartment) in the decidua and placenta. The prevalent activation of Tregs that our analysis points to in those organs may be associated with the requirement for immune tolerance during pregnancy (**Supplementary Fig. 12B right**).

Comparing Tregs that came from healthy vs. pathological samples, we detected a general shift towards activated Tregs in diseased individuals (**Supplementary Fig. 12B left**). This trend is driven, to some extent, by the milieu of cancer samples that are included in the Census, which tend to have high proportions of activated Tregs within the Treg compartment (**Fig. 5F**). Indeed, our activated Treg population expresses higher levels of *TNFRSF4*, *TNFRSF9*, *TNFRSF18*, *ICOS*, *CTLA4* and *LAG3* while our memory Treg population expresses higher levels of *PI16*, *ITGB1* and *CCR10* (**Supplementary Fig. 13 and Fig. 5E),** which lines up well with the signature of tumor-infiltrating Tregs in <u>(Kim et al. 2021)</u>. Interestingly, while most cancers tend to be enriched with the activated Treg state, we found a group of cells from a breast cancer study (labeled as cancer cells in their meta-data) that were an exception. Following up on this part of the dataset, however, we found that these cells were not collected from the cancer tissue but from the contralateral breast during cancer-related mastectomy <u>(Kumar et al. 2023)</u>. Indeed, a separate study has identified these activated Tregs in breast cancer expressing several of the marker genes identified above <u>(Plitas et al. 2016)</u>. Put together, this analysis serves as an example of how a cellular population of interest (here Tregs) can be identified in the Census, and how the myriad conditions and pathologies that are included in this collection can be leveraged, e.g., for finding the pathological conditions or tissue locations in which they are prevalent.

## DISCUSSION

Given the growing corpus of single-cell omics datasets (both individual studies and atlas-level efforts), techniques for transfer learning are becoming pivotal in enabling studies of both new and revisited datasets. Despite the promise of these techniques, their use has been limited owing to two major reasons. First, the use of large reference datasets can be prohibitive both in terms of the required compute resources (e.g. to access and process the data) and expertise (e.g. for integrating it with query data). Second, there is a need for an appropriate platform to facilitate communication between data providers and consumers and to provide the infrastructure for quality control, access and downstream analysis. Scvi-hub provides a way to alleviate these by establishing a platform for sharing and reusing single-cell omics data, by not only making the data itself available but also making available models that have been trained on that data. The sharing of models helps lower the need for technical expertise and expensive compute resources (e.g., to process atlas-scale references and integrate them with query data) and the respective API was designed to facilitate a wide array of analyzes.

The decentralized nature of scvi-hub, relying in part on the properties of the platform it is hosted on (Hugging Face) as well as supporting AWS s3 buckets, enables community access through friendly and easy-to-use interfaces. As such, we envision scvi-hub to serve several types of users. First, we expect it to become a platform for individual researchers who wish to not only share their data post-publication, but also make their analyzes accessible and reproducible. Second, we expect it to be used in efforts for generating tissue atlases as well as large scale single-cell census datasets, as a way of facilitating advanced use of these atlases (beyond exploratory analysis). Third, we expect the research community to leverage the models in scvi-hub as an actionable resource for an array of use-cases, from annotation of new scRNA-seq samples to deconvolution of ST samples. To demonstrate this, we presented several case studies showing that incorporating external references can improve and enrich the analysis of individual datasets and provide novel insights into disease mechanisms and the pertaining effects on cell type composition and gene expression.

The model-centric approach of scvi-hub enables representation of large reference datasets in a minified format, which enables access with limited memory resources or download bandwidth and thereby accelerates access to those valuable resources. It is our hope that this will help democratize single-cell data analysis and expand it to communities with low compute resources and/or enable work in scenarios where such resources are scarce. Validating findings in the original expression space, however, is key. Scvi-hub can serve as a central gateway to data repositories, like CELLxGENE Discover (CZI Single-Cell Biology Program et al. 2023), which allows users to access selected portions of very large datasets (e.g. genes or subpopulations highlighted by the model-based analysis) for close inspection. We highlight this capability by reanalyzing an existing CAR T datasets using the knowledge accumulated over the full data repository and identifying novel cell populations that are infused to patients and might have an impact on side effects or efficiency of treatment. Scvi-hub can harmonize annotations across the diverse sets of datasets in those repositories and thereby provide novel insights into shared features between diseases. We highlight this by identifying a TREG population shared across all types of cancer well represented in the current repository. Identifying those shared features, will in the future help to refine our understanding of cellular subtypes. It also became obvious that our current way of structuring metadata is limited as e.g. *in vitro* stimulated cells are annotated as healthy cells or cells from a healthy tissue are labeled by a disease existing in a different tissue (like contralateral breast during breast cancer). Efforts like HuBMAP work to harmonize these annotations <u>(Human BioMolecular Atlas Program 2023)</u>, this will be critical in the future to further increase the value of these cross-study analyzes.

An important part of the interface between data providers and consumers is the ability to criticize and reach informed decisions about the merit of a given reference dataset or model for an application of interest. To that end, we developed a new suite scvi-criticism, that can serve to validate models prior to upload as well as for evaluating how well a query dataset fits a reference model. We believe both are essential for effective transfer learning.

The development of scvi-hub aims to foster a model-driven paradigm in the single-cell data analysis community – one where models are easy to find, access, develop, and share, and can be efficiently leveraged to analyze various aspects of new and existing datasets. We expect it to become a growing resource, catering for new types of analyzes, use-cases, and data modalities.

## METHODS

### HubModel

Scvi-hub is implemented as a lightweight submodule within scvi-tools (*scvi.hub*), which provides an API for uploading and downloading pretrained models to the Hugging Face Model Hub. To do so, scvi-hub uses the *huggingface_hub* Python API. The main construct within *scvi.hub* is the *scvi.hub.HubModel* class, which represents a pretrained scvi-tools model hosted on the Hugging Face Model Hub. An instance of this class has properties that can be used to load the model (*HubModel.model*) and data (*HubModel.adata*) into memory on demand (i.e. only when/if the property is invoked). To help with new model creation and upload, we also provide a *scvi.hub.HubModelCardHelper* class – which can be used to auto-generate a template Model Card for a new model to be uploaded to the hub – and a *scvi.hub.HubMetadata* class which encapsulates metadata required to be uploaded with the model to the hub. Model versioning is built into scvi-hub via usage of Hugging Face as our storage platform, since Hugging Face Hub Models are backed by git. Hub users can thus request a specific version of the model at download time via the *revision* argument to the *pull_from_huggingface_hub* function. We additionally support Hub Models uploaded to an AWS s3 bucket using *scvi.hub.HubModel.pull_from_s3*. This allows interacting with models that are stored privately inside company buckets or shared with the community without the requirement to register to HuggingFace.

### Data minification

We allow to store the latent posterior parameters (variance and mean in the latent space for each cell). This allows for faster generation of estimated gene expression as the inference function of scvi-tools models aren’t executed and instead the stored latent posteriors are used for the generative parts of the models. In addition, we can reduce the dataset size by removing all count data in the AnnData object. This is performed by setting all expression values to zero and storing count objects as a sparse matrix (therefore zero entries in the sparse matrix). We additionally allow storing latent posteriors without removing the associated count data. A few functions in scvi-tools rely on computing the reconstruction loss and therefore require access to the full count data. These functions are not executable after data minification. All code used throughout the manuscript is compatible with data minification.

### Evaluation using scvi.criticism

Scvi.criticism is a new submodule in scvi-tools that can be used to efficiently compute model evaluation metrics for scvi-tools models. Models must inherit from the scvi’s *BaseModelClass* class and implement the *posterior_predictive_sample* method, which samples from the generative distribution used by the scvi-tools model such as a negative binomial distribution. The main entry point for using scvi.criticism is the *PPC* class, which computes and stores various metrics for the provided collection of models. The *PPC* class can be initialized with one or a collection of models (for instance for model comparison), the raw counts, and a host of configuration options (such as number of posterior predictive samples to compute). It internally computes and stores a user-provided number of posterior predictive samples for each model (samples from the generative function of the model with user-provided library size).

A crucial aspect of this package is integrated support for 3D sparse arrays. As the posterior predictive samples are *n_cells* x *n_features* x *n_samples* data cubes, their size grows linearly with the size of the dataset and the number of posterior predictive samples. We use the sparse (https://github.com/pydata/sparse) and xarray (https://github.com/pydata/xarray) Python packages to store the raw counts and posterior predictive samples in sparse format in memory, and only hydrate the data one batch at a time (this logic is implemented in the scvi-tools package). Supporting sparse arrays in scvi.criticism enables efficient evaluation of models trained on large-scale datasets.

Once initialized, the *PPC* class can be used to compute various metrics on the per-model posterior predictive samples. The class will store a collection of metrics keyed by metric names where each entry is a Pandas DataFrame holding results for all models that the class was initialized with.

We implemented the coefficient of variation, which is defined as the standard deviation over genes or cells divided by the mean over the same axis. Respectively, the user decides whether the metric is summarized over the different genes or different cells.

To generate **Fig. 2D**, we instantiated the *PPC* class with the HLCA raw counts data, the pre-trained scANVI model, and a value of 2 for the number of posterior predictive samples. The top plot shows the coefficient of variation metric computed across the “features” dimension, using the *coefficient_of_variation* method of the *PPC* class. We generated 2D histograms shown along with the best fit and identity lines. The method also prints out correlation measures (such as R2, Pearson and Spearman correlations, and mean absolute error) between raw and approximated results. Results can be stored as an *obs* column in *AnnData*. For **Fig. 2D** bottom, we computed the log-ratio of coefficient of variation between the scANVI model and raw (without adding any pseudocounts) and displayed the corresponding values overlaid on UMAP.

We implemented the differential expression (DE) metric. For each of the posterior predictive samples differentially expressed genes are computed separately. Samples are first library size normalized and log1p-transformed. We execute *scanpy.tl.rank_genes_groups* with a user-provided column for the cell-types and the respective method for computation, *t-test* by default. For each sample, we afterwards compute the F1 score of the top 100 overlapping genes, the mean absolute error (MAE), Pearson correlation and Spearman correlation between all estimated log2-fold changes and the AUC under the ROC curve and the average precision score for all genes with a significant p-value computed on the raw expression (significance threshold - *pvalue_thresh* by default 0.001). **Fig. 2E** shows the differential expression metric computed using the *differential_expression* method of the *PPC* class, “ann_level_3” key for cell types and a *pvalue_threshold* of 0.2. We used the *plot_diff_exp* method of the *PPCPlot* class to generate the box plot shown. In **Fig. 2F**, we subsetted the dotplots to the immune cell subtypes, i.e. those with a cell type label in the following list: “B cell lineage”, “Dendritic cells”, “Macrophages”,“Mast cells”, “Monocytes”, “T cell lineage” and display the log-2 fold-changes of the respective top 2 marker genes. To generate the dotplots, we used one sample of the posterior predictive samples

In **Supplementary Fig. 3**, we show the results of *coefficient_of_variation* (**a**) and *differential_expression* metrics (**b**) computed on a subset of the heart cell atlas dataset (Litviňuková et al. 2020). We preprocessed the dataset as presented in this tutorial (https://docs.scvi-tools.org/en/stable/tutorials/notebooks/api_overview.html), then trained three models on the preprocessed data as follows: One model was trained with the default max number of epochs (400) and latent dimensions (10). The second model was trained with only 5 epochs (and the default number of latent dimensions) and the third model was trained with only 2 latent dimensions (and the default max number of epochs). (**a)** was generated in the same way as the 2D histogram shown in **Fig. 2E**. (**b)** was generated in a similar fashion to the **Fig. 2E**.

In **Supplementary Fig. 4**, we downloaded all epithelial cells from Tabula Sapiens (https://cellxgene.cziscience.com/collections/e5f58829-1a66-40b5-a624-9046778e74f5) and used those cells as query cells for the pre-trained HLCA scANVI model. We set the batch_key to the respective tissue to learn independent transfer mappings for each organ. We computed the CV over cells separately for each tissue and plotted the results similar to the top plot in **Fig. 2D**.

### Reference-based analysis

For this analysis, we used the Human Lung Cell Atlas (HLCA) (Sikkema et al. 2022) dataset and their pre-trained scANVI model. We used *cellxgene_census* to download the dataset (implemented in scvi-tools) and used for all analysis the raw, unnormalized counts. We used scvi-hub to download the model from the Hugging Face Model Hub. The author-provided UMAP embeddings are used and Scanpy is used for downstream analysis (Wolf, Angerer, and Theis 2018). The celltype UMAP is representative of the “ann_level_3” annotations. To generate **Fig. 2C**, we used the minified data from Hugging Face, then generated expression values of the genes displayed using a single sample from the posterior predictive checks. For the displayed raw expression as well as model-generated values, we computed library-size normalized and log1p-transformed values.

### Transfer learning analyses

For all models trained using transfer learning, we used following parameters *surgery_epochs=500* (200 for emphysema dataset), *early_stopping=True*, *early_stopping_monitor=’elbo_train’*, *early_stopping_patience=10*, *early_stopping_min_delta =0.001*, *weight_decay =0.0*. For Tabula sapiens epithelial cells, each tissue was treated as a separate batch inside scANVI to allow comparisons between the various organs, while for both the emphysema data set and the cross-organ immune data set all cells were treated as a single batch.

We used the same HLCA reference model as in Fig. 2 as the reference data, and the data set from human lung emphysema as the query data. We used the scArches functionality implemented in scvi-tools to prepare and train a model on the query dataset. We then used the *get_latent_representation* method of the concatenated reference and query dataset to retrieve coordinates of the query data embeddings in the joint reference/query latent space.

To generate **Fig. 3A**, we computed Scanpy’s nearest neighbors in the combined latent space using *n_neighbors=30* and UMAP using *min_dist=0.3* in the RAPIDS implementation.

We transferred labels to the query dataset by first learning a nearest neighbors index on the latent space of the reference atlas, using the *NNDescent* class of the pynndescent package, and then using the index to compute a nearest neighbor graph for the query dataset. We then used this graph to assign to each cell in the query dataset, a predicted cell type based on the reference dataset, along with a prediction uncertainty (we used the “ann_level_3” for the cell type annotation in the reference dataset). To this end, we converted nearest neighbor distances to affinities, and weighted the predictions using these affinities (this follows the approach used in the HLCA). For the emphysema dataset, we only use predictions for the confusion matrices with an uncertainty below 0.4 (4.8% of cells filtered out for emphysema dataset). For transferring labels from the query dataset to the reference dataset, the same function was used but the role of reference and query dataset were replaced. We used an uncertainty threshold of 0.2 (52.1% of cells filtered out of extended HLCA dataset) for the confusion matrix.

In **Fig. 3C**, we display summarized cell-type labels after recomputing UMAP using Scanpy with *n_neighbors* set to default. In **Supplementary Fig. 5**, we display all labels predicted by label transfer and their confusion matrix with the original labels (normalized by columns). In **Fig. 3D**, we compute differential abundance using Milo between healthy and diseased cells from the query dataset using 100 neighbors in the latent embedding space, treating donor_id as the sample columns and not correcting for any covariates. Results are displayed for an FDR < 0.1.

In **Fig. 3E**, to compute differentially expressed genes, we set *idx1* to all fibroblast from diseased individuals and *idx2* to fibroblast from healthy individuals. Fibroblasts were identified as all cells labeled as *Distal fibroblast*, *Proximal fibroblast* or *Peribronchial fibroblast* in the original publication. The *differential_expression* function of the HLCA model was used filtering for outlier cells and correcting the batch. Of note, *weighting=’Importance’* requires access to the full expression values of the reference dataset and was deactivated here. Genes were selected to be expressed with an estimated mean (scale in scvi-tools) of >1e-4 in either of both groups. For the volcano plot decoupler-py was used highlighting the top 20 genes. The mean log-2 fold-changes are displayed on the x-axis and *proba_not_de* on the y-axis. In **Supplementary Fig. 6, we first display UMAP embeddings colored by the donor and disease status to corroborate the differential abundance analysis and display DE genes overlaid on UMAP to show that most genes are expressed across fibroblasts**. In Fig. 3F, violin plots were created using Scanpy on library-size normalized and log1p-transformed raw gene expression values.

In **Supplementary Fig. 7, for PyDESeq2, we used decoupler**-py and filtered out all genes with *min_count=50* and *min_total_count=150* grouped by disease status. Pseudobulks were computed per donor. All genes were used in the top panel, while for the bottom run we subsetted all genes to the ones incorporated in the reference model before running PyDESeq2.

In **Supplementary Fig. 8, results of DestVI on prostate tissue are displayed**. We accessed the prostate tissue single-cell model from Hugging Face and ran deconvolution For **Fig. 3G**, we downloaded all T cells and innate lymphoid cells from CELLxGENE Discover and subset this object to all cells originating from the lung. Analysis was performed as described above. Labels were infused to the extended HLCA dataset and confusion matrix is displayed in **Fig. 3H** similar to **Supplementary Fig. 5**. For **Fig. 3I**, marker genes were hand-selected and expression is displayed using library-size normalized and log1p raw expression values. For **Fig. 3J**, we subset to all cells labeled as Tem/Temra_CD8 and used PyDESeq2 with same settings as above for **Supplementary Fig. 7**. **Supplementary Fig. 9, contains recomputed UMAPs on query and reference T cells**. The label-infused cell-types, the original query annotation and the source dataset are displayed as well as the same marker genes of regulatory T cells as in **Fig. 2C**.

### CELLxGENE census reference model

We used a custom trained model trained on the CELLxGENE release and released afterwards embedding via CZI with similar quality. For our training run, we subset the CELLxGENE census object to all protein coding genes using the PyBiomart interface in Scanpy as well as subsetting to features included for more than 30 million cells inside the census release (LTS 12/2023). We identified 6000 highly variable genes using the census highly_variable_gene function and using suspension_type and assay as batch keys. We afterwards took the intersection of those highly variable genes and protein coding genes. We trained the scVI model using following custom parameters (optimized by visual inspection of multiple training runs): 2 layers, no encoding of covariates, 50 latent dimensions, 512 hidden nodes, batch size of 1024, early stopping enabled with monitoring elbo_validation, n_epochs_kl_warmup of 5 epochs, learning rate of 1e-4, max_epochs 20, batch key as concatenation of suspension_type, dataset_id, donor_id and assay). We updated the parameters of the scVI model published by CZI afterwards and validated that it yielded similar results to our custom trained model. In the future, CZI will release new trained models to integrate novel datasets for every LTS model (expected every 6 months). For the CZI released model, genes are not subset to protein coding genes, *n_epochs_kl_warmup* was increased to 20 epochs, *max_epochs* was increased to 100 epochs and *early_stopping* was disabled. We made these changes to have reliable training runs for future releases of the census datasets.

We downloaded the CAR T cell study from CELLxGENE and used transfer learning to embed it running for 1000 surgery_epochs with early_stopping enabled and using a batch size of 4096 to accelerate training. The model was stopped early after 556 epochs and it took 1 hour and 50 minutes to complete training. A subset of 10% of the primary cells in census and the CAR T cells were concatenated and a joint PyNNDescent index was built and labels propagated to all cells from the census dataset. For Figure 4 B and C, a random subset of 10% of the original cells was used and UMAP was computed using default parameters within the RAPIDS framework in Scanpy. To reannotate the query dataset, a PyNNDescent index was built on cells from the cross-tissue immune cell atlas and query cells were annotated similar to figure 3. Differentially expressed genes were identified using the scvi-tools differential expression function and we display the top 3 markers with lowest p-value and an estimated scale of the negative binomial distribution above 1e-4. Louvain clustering was performed using 50 nearest neighbors and default resolution. Milo was performed using 100 nearest neighbors in scVI latent space, treating donor_id as the sample column and not correcting for any covariates. Results are displayed for neighborhoods with an FDR < 10%. Differential abundance for CRS state was compared between no CRS and grade 3 and 4 CRS. Grade 1 and 2 CRS contained diverse cells. To display marker genes of the Louvain clusters, we hand selected genes and displayed them using the dotplot plotting function in Scanpy.

### Reanalysis of census dataset

The same trained model was used as for Figure 4. To build a neighborhood index, we concatenated the cross tissue immune cell atlas and a similar number of randomly selected cells from the CELLxGENE census. The other cells were labeled as “census” and built a PyNNDescent index on those cells. We set the number of neighbors to 50 cells. We display hand-selected marker genes in **Supplementary Fig. 12C**. We subset all cells to those cells labeled “regulatory T cells’’. We downloaded the raw expression values of these cells. An scVI model was trained using the same gene subset as for the large model with default parameters except using 2 layers and *n_epochs_kl_warmup* of 50 epochs. UMAP was computed using 20 neighbors and a *min_dist* of 0.3 in the RAPIDS framework. Subcluster markers were hand-selected. We excluded all clusters with a low expression of *FOXP3* from further analysis. Barplots in **Figure 5 D and F** were generated with matplotlib. In **Figure 5D**, all bars together add up to a width of 1, while in **Figure 5F** relative proportions per disease category are displayed. Pseudobulk differential expression analysis was performed using PyDESeq2 setting min_cells to 5 and min_counts to 1000 with the concatenation of dataset_id, donor_id, disease and tissue as the sample column and the identified Treg subclusters as the group column. We filtered out genes with a min_total_count of 15 and 2 counts per sex category. Including disease as covariate drastically increased runtime and we stopped the algorithm after an hour. Therefore, we excluded covariates from analysis and only used the Treg subclusters in the formula. We display the top 50 genes based on the adjusted P value between ‘activated-TREG’ and ‘memory-TREG’.

## Availability of data and materials

We used the Human Lung Cell Atlas (HLCA) dataset in our analysis, which can be found here (https://cellxgene.cziscience.com/e/066943a2-fdac-4b29-b348-40cede398e4e.cxg/) and their pre-trained scANVI model which can be found here (https://zenodo.org/record/6337966/files/HLCA_reference_model.zip). We provide tutorials for data minification (https://docs.scvi-tools.org/en/stable/tutorials/notebooks/minification.html), as well as how to implement this feature for newly-developed latent variable models.

As additional single-cell datasets, we used several datasets from CELLxGENE Discover. Namely we accessed Tabula sapiens data at (https://cellxgene.cziscience.com/e/53d208b0-2cfd-4366-9866-c3c6114081bc.cxg/) and respectively the epithelial cells at (https://cellxgene.cziscience.com/e/97a17473-e2b1-4f31-a544-44a60773e2dd.cxg/). The emphysema dataset was downloaded from (https://cellxgene.cziscience.com/collections/03cdc7f4-bd08-49d0-a395-4487c0e5a168). All files were downloaded. The cell-type information of AT2 cells was used and all three separate datasets were concatenated and treated as one dataset for query analysis. The cross-tissue immune-cell dataset was downloaded from (https://cellxgene.cziscience.com/e/ae29ebd0-1973-40a4-a6af-d15a5f77a80f.cxg/). The Heart Cell Atlas was downloaded from (https://cellxgene.cziscience.com/collections/b52eb423-5d0d-4645-b217-e1c6d38b2e72). The Visium dataset for the prostate was downloaded from 10X (https://cf.10xgenomics.com/samples/spatial-exp/2.0.0/Visium_FFPE_Human_Prostate_IF/Visium_FFPE_Human_Prostate_IF_spatial.tar.gz).

All datasets were accessed and used in the version of December 1st, 2023 and all files can be shared upon contact with the corresponding author.

A GitHub repository with extensive code to reproduce all figures is provided at https://github.com/YosefLab/scvi-hub-reproducibility.

## Acknowledgments

We thank Gonzalo Benegas for the suggestion to host our pretrained models on the Hugging Face Model Hub. We thank Pierre Boyeau and members of Yosef laboratory for general feedback. We also acknowledge ChatGPT in generating code. We thank Emanuele Bezzi, Bruce Martin, Andrew Tolopko, and Pablo Garcia-Nieto from the Chan Zuckerberg Initiative for their support on making available trained scVI models and their associate latent spaces for all data in CELLxGENE Discover Census (https://cellxgene.cziscience.com).

## Author contributions

C.E. and V.V.P.A. contributed equally. C.E., A.G., and N.Y. conceptualized the study. V.V.P.A. implemented scVI-hub with the help of A.G., C.E. and M.K. C.E. performed the validation experiments with inputs from N.Y., V.V.P.A. and A.G. N.Y., A.S. supervised the work. C.E.,

V.V.P.A. and N.Y. wrote the manuscript with input from A.G., A.S., M.K.

