## Supplementary Figures for "Scvi-hub: an actionable repository for model-driven single cell analysis"

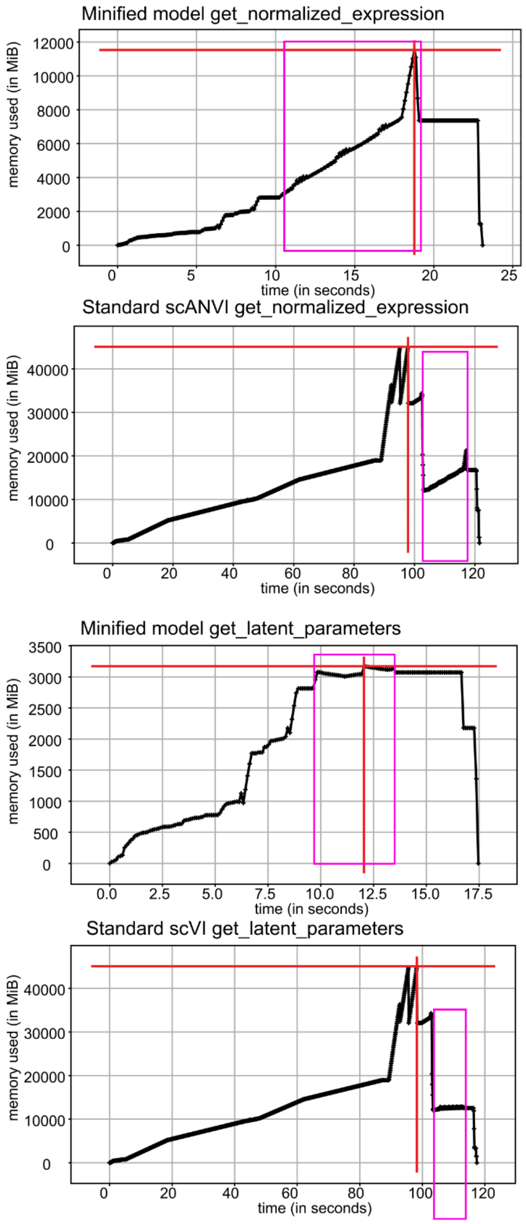


**Supplementary Figure 1: Hardware improvement using minified data.** Memprofiler was used to compare performance of *model.get_normalized_expression*, which requires encoding and decoding of cells and stores a dense copy of a cell-by-gene expression matrix and *model.get_latent_parameters*, which requires encoding cells. The scripts include loading the minified or the full AnnData, preparing AnnData for use inside scvi-tools and calling the respective function. In pink boxes the actual time to compute the scvi-tools function is provided. We set the minibatch size to 16000 to reach maximum speed for all functions. The largest time for non-minified data is spent to actually read the dataset and memory also spikes here at 43 GB. Memory usage is reduced when subsetting the genes used for model training to 13 GB which is four times more than the minified dataset (3 GB). The normalized gene expression requires 9 GB of RAM totalling at 12 GB RAM for ~600k cells for minified data. Computing the normalized expression is roughly 50% faster using minified data (7 seconds versus 15 seconds).

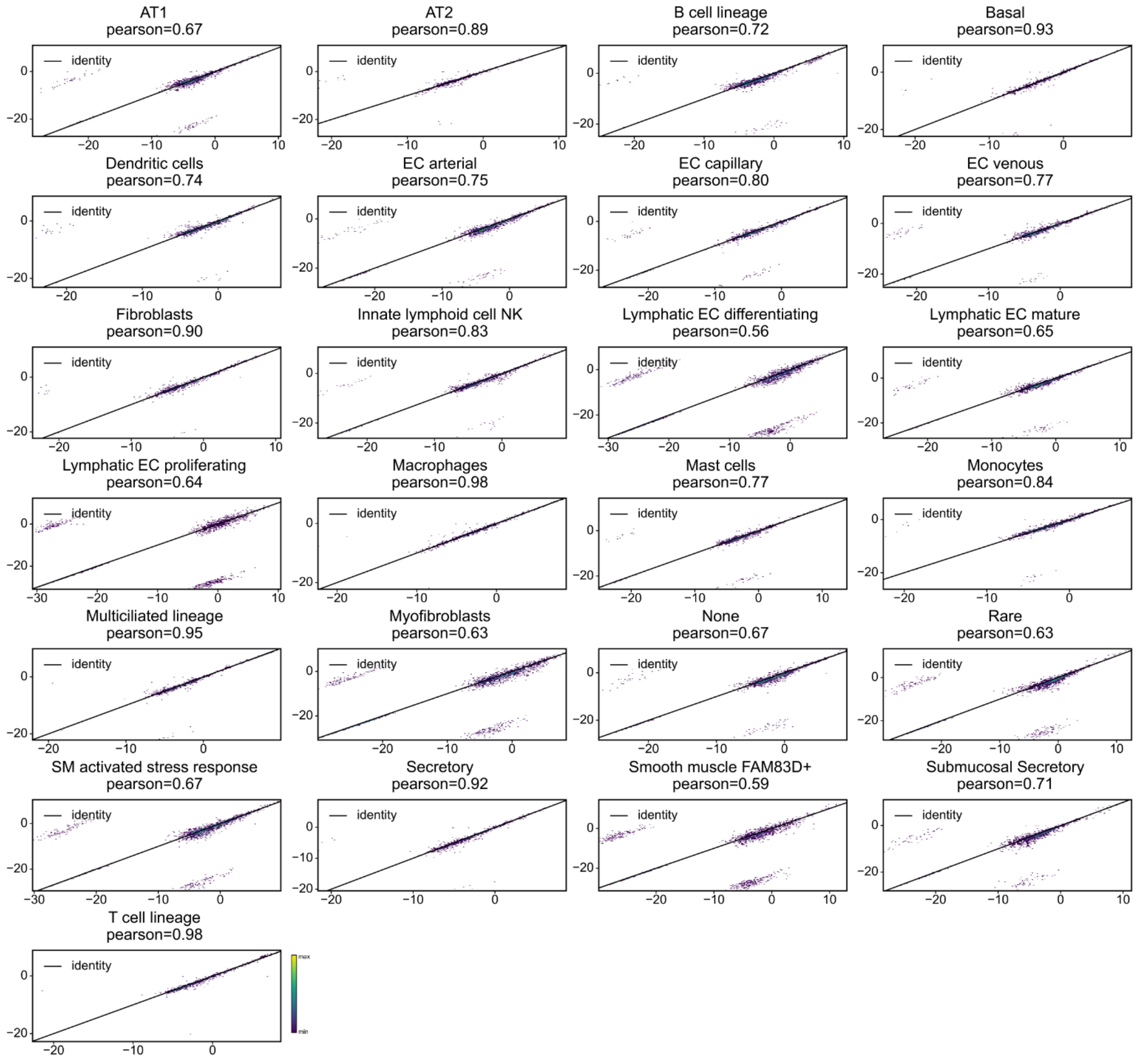


**Supplementary Figure 2: Highlighting per cell-type differential expression based metric.** The differential expression based PPC metric is displayed for each cell-type separately. We find a Pearson correlation coefficient of above 0.6 for each cell-type. We detect a lower correlation coefficient for cell-types with only a few cells. While all upregulated genes have a strong positive correlation. We find for some genes a highly negative log-2 fold-change estimate. This is due to numeric issues and those genes are almost not expressed in the respective cell-type leading to prediction of a strong downregulation of these genes.

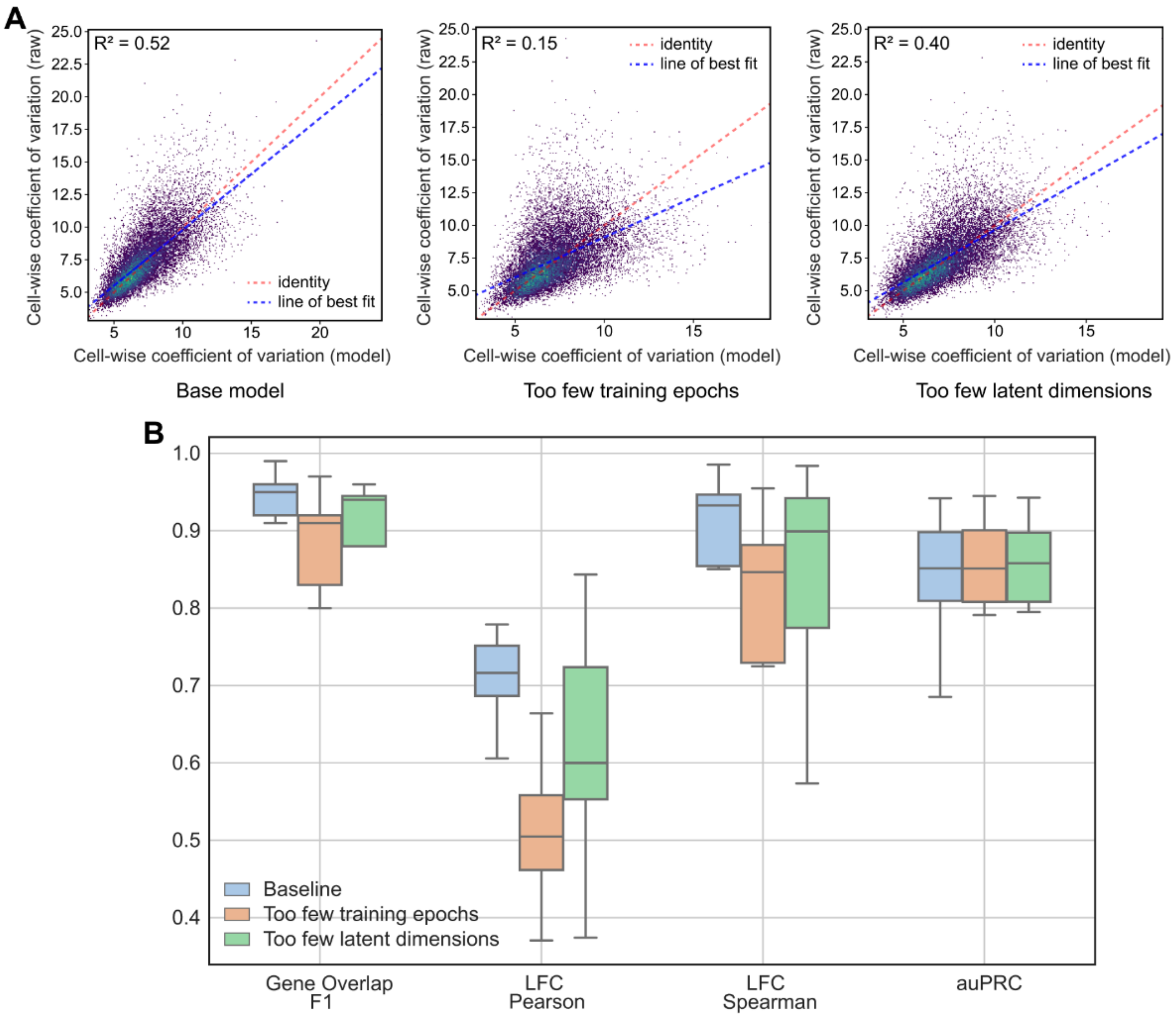


**Supplementary Figure 3: Scvi-criticism detects poorly trained models. (A)** Cell-wise coefficient of variation for 3 different models trained on the Heart Cell Atlas data: the left model is the base model trained for 400 epochs with 10 latent dimensions, the middle model is trained for 5 epochs with 10 latent dimensions, the right model is trained for 400 epochs with 2 latent dimensions. The Pearson correlation coefficient is lower for both models trained with corrupted parameters. **(B)** Differential expression based metric with F1-score of top-100 genes between the different celltypes, second and third Pearson and Spearman correlation coefficient between all estimated log-2 fold-changes, fourth area under the precision-recall curve (auPRC). All metrics despite auPRC display a reduced performance for those corrupted models compared to the default model.

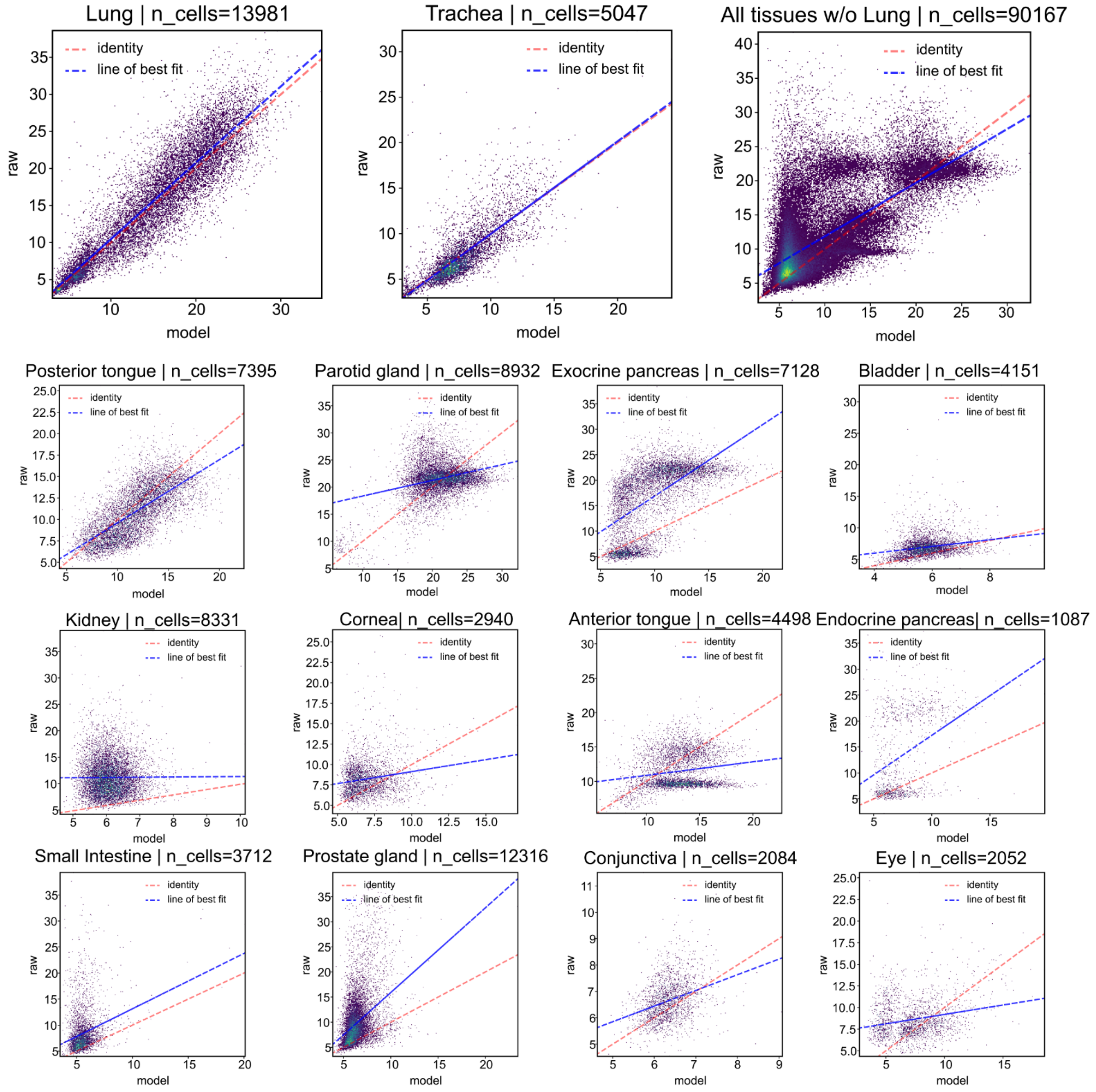


**Supplementary Figure 4: Scvi-criticism detects query data not suited for a reference model.** We trained a query model based on the HLCA reference model using all epithelial cells from Tabula sapiens. Coefficient of variation is displayed separately for each organ. For lung, trachea and posterior tongue (slightly worse) we find a high correlation of raw and model coefficient of variation. For all other tissues (top right) we find a much worse capturing of the CV by the trained model. For bladder, parotid gland, kidney, corneal, anterior tongue and eye there's no correlation between the model estimated CV and the raw data CV with the model estimated CV overestimating the raw data CV. For other organs like prostate gland and exocrine and endocrine pancreas the raw CV is in general higher than the model estimated CV with a low linear dependency. Of note, highly variable genes were selected initially based on lung tissue and this causes the small CV for cornea and eye epithelial cells.

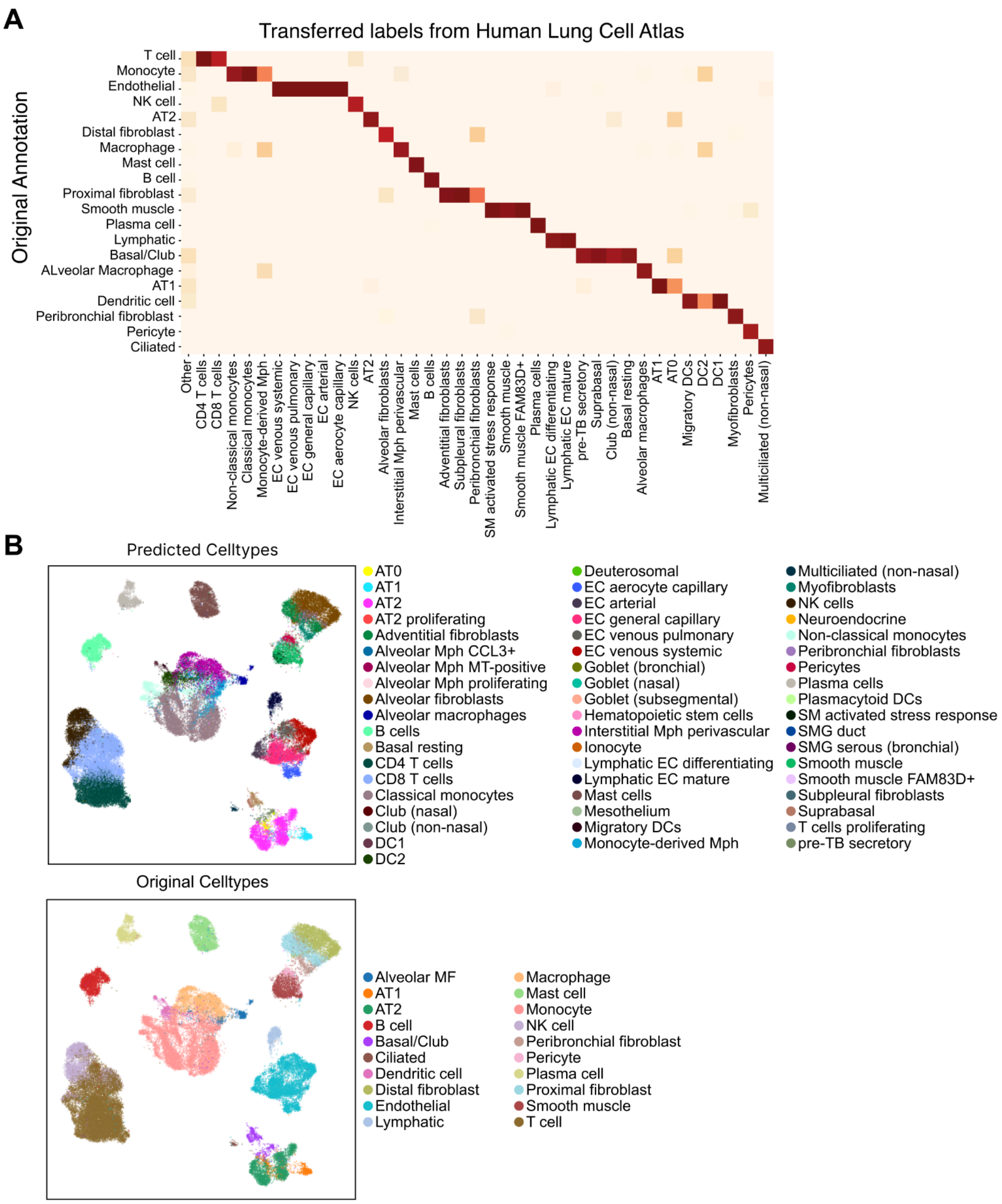


**Supplementary Figure 5: Cell-type label transfer for emphysema dataset highlights concordance between annotations.** HLCA labels were transferred to the emphysema dataset. **(A)** Confusion matrix for all cells with high certainty for the transferred label (<10% of cells excluded with a low certainty of the transferred label). On the y-axis it shows the original annotation and on the x-axis the transferred label. There is overall agreement in the coarse tissue group. For several cell-types the transferred labels have a higher resolution. All cell-types predicted less than 10 times are summarized in the other column. Values are normalized per column **(B)** (top) UMAP as in Figure 2C (with cells colored by their transferred labels), whereas in Figure 2C cell-types were summarized for visualization purposes. (bottom) Cells organized in the same UMAP and colored by their original labels. In comparison, it becomes evident that transferred labels yield a higher granularity of labels while both labeling schemes are concordant.

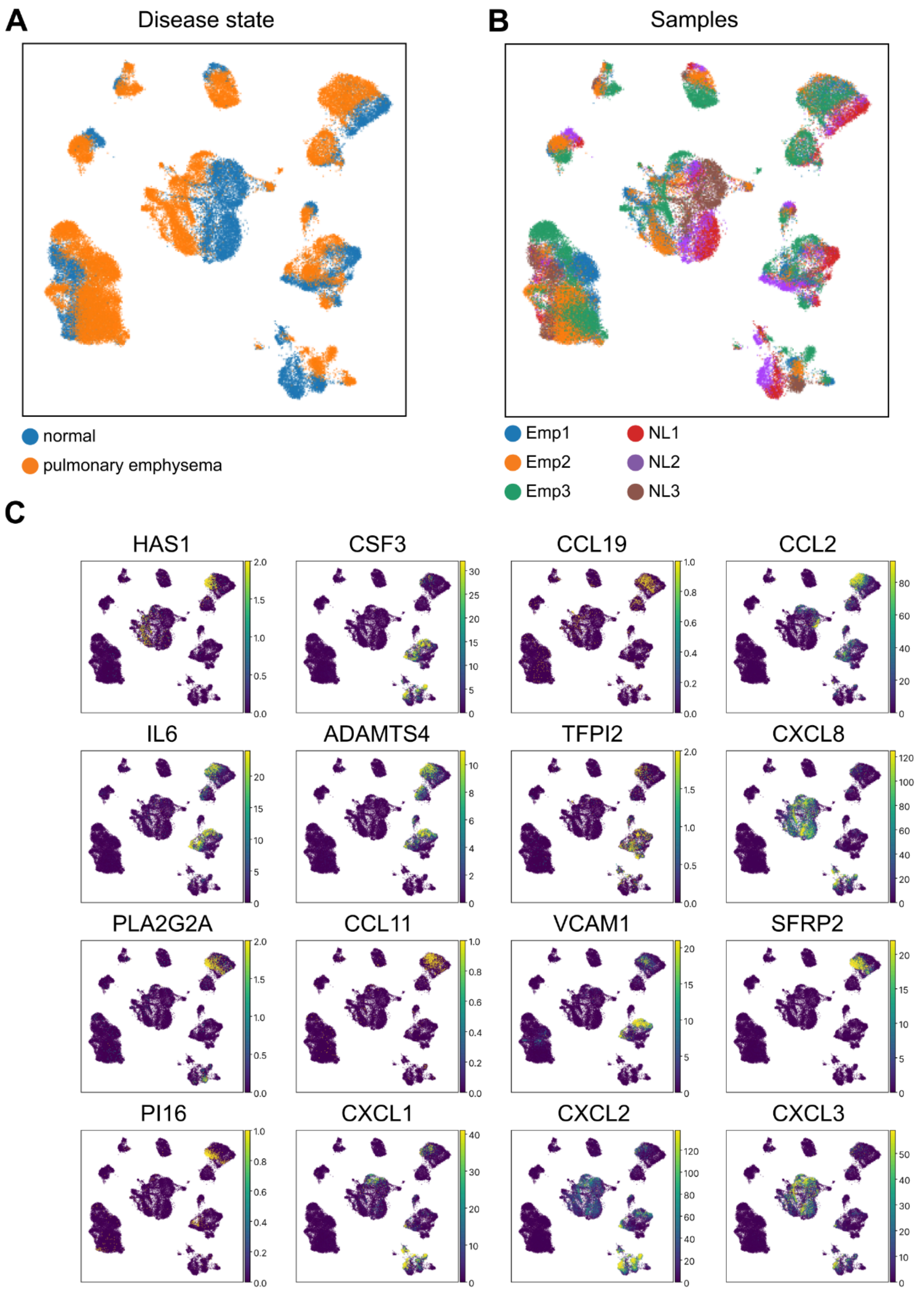


**Supplementary Figure 6: Differentially expressed genes between healthy and diseased fibroblasts. (A)** UMAP embedding colored by healthy control cells (*normal*) and diseased state (*pulmonary emphysema*) underlines the results of the differential abundance analysis in Figure 3d. **(B)** UMAP embedding colored by original donor ID. Highlights strong intra-individual differences for myeloid cells and T cells and low intra-individual differences for fibroblasts. Due to the consistency in signal, fibroblasts were used for downstream analysis. **(C)** Cellular expression pattern for all genes discussed in the main figure highlights that indeed these genes are expressed in fibroblasts in a relevant amount and the region, which is more abundant in cells from diseased individuals indeed expresses higher levels of these genes. Expression values from raw counts after library-size normalization and log1p-transformation are displayed.

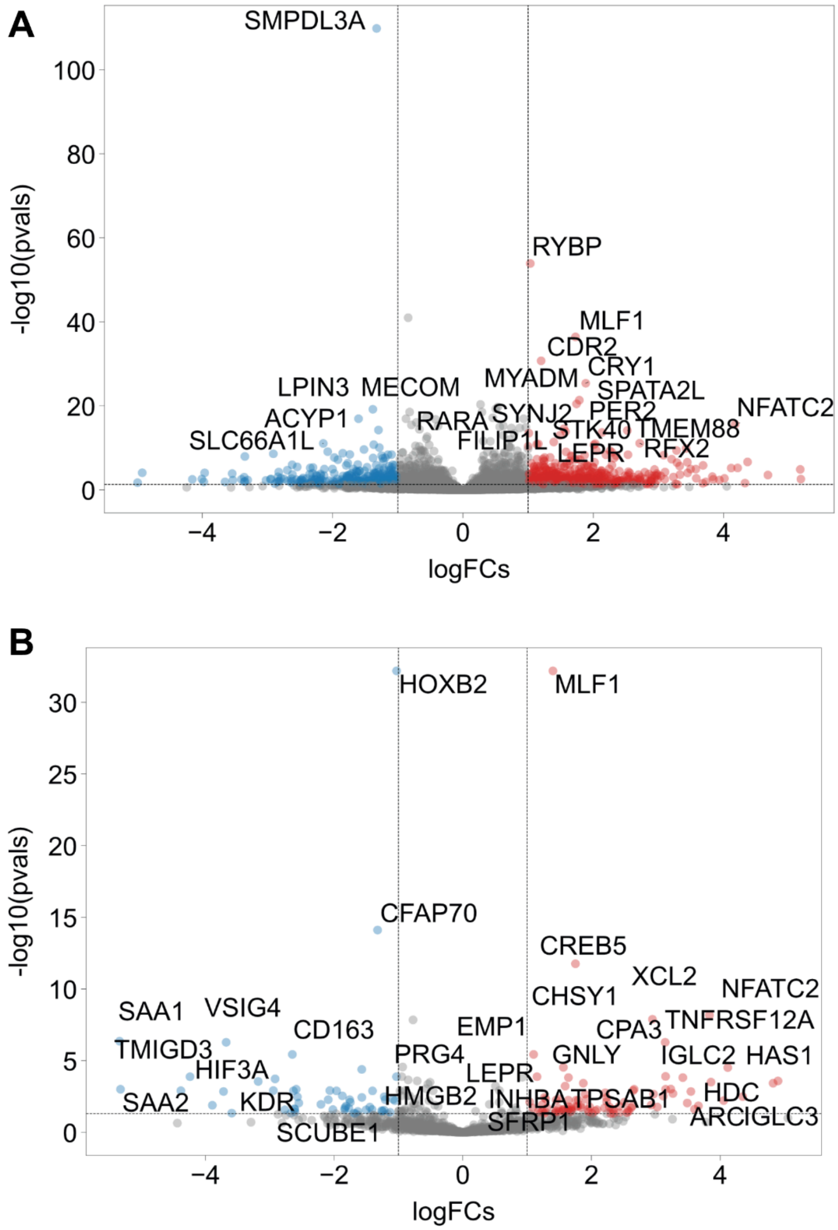


**Supplementary Figure 7: Pseudo-bulk differential expression test does not reveal similarly relevant differentially expressed genes. (A).** Pseudo-bulk DE analysis considering all genes after filtering for lowly expressed ones. Genes listed as up-regulated are described to be linked with tumorigenesis. These genes have no specific role in fibroblasts and their role in emphysema remains unclear. **(B)** We additionally computed differentially expressed genes after subsetting to the genes used to train the HLCA model (i.e., the genes that were considered in the model-based DE in **Fig. 3**). Differentially expressed genes are enriched in canonical marker genes of other cell-types (GNLY, XCL2 lymphoid cells; TPSAB1 mast cells; CD163, VSIG4, TMIGD3 in macrophages). They also include genes associated with P53 signaling such as CREB5, NFATC2, CPA3 and SFRP1, which are expected to be upregulated in smoking individuals (. These genes were also identified with the model-based analysis.

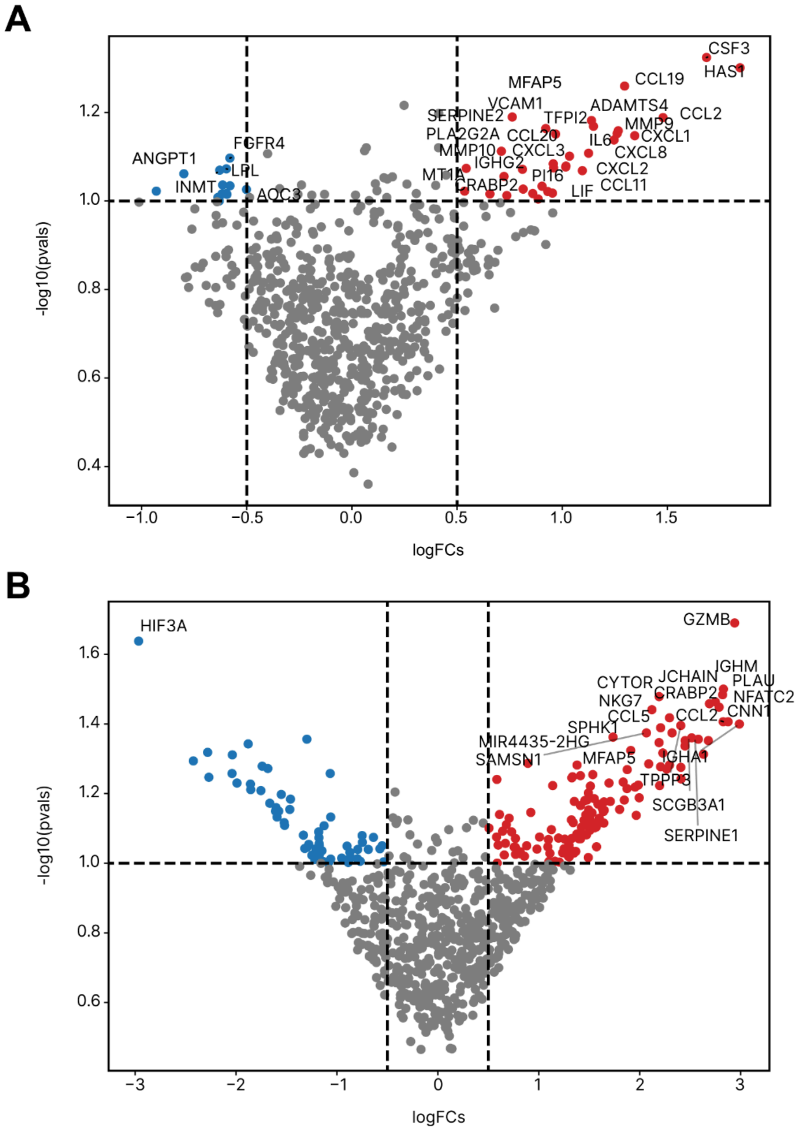


**Supplementary Figure 8: Ablation study for differential expression function in scvi-tools.** Both figures highlight differentially expressed genes with exactly the same settings as in **Fig. 3E**. **(A)** A query model was trained using the same setting as in **Fig. 3E**. The batch_key in scANVI was set to the donor ID instead of providing a single batch_key to the query model. In the DE function, we are not using transform_batch setting and instead generate the estimated expression using the original donor IDs. **(B)** A query model was trained using the same settings as in **Fig. 3E**. During *scvi.model.SCANVI.load_query_data*, we changed *unfrozen=True*, meaning that every component in the reference model is updated based on the query data. Bottom plots displays much more differentially expressed genes, with a high number of genes actually expressed by other cell-types than fibroblasts (mainly CD8 T cells and plasma cells).

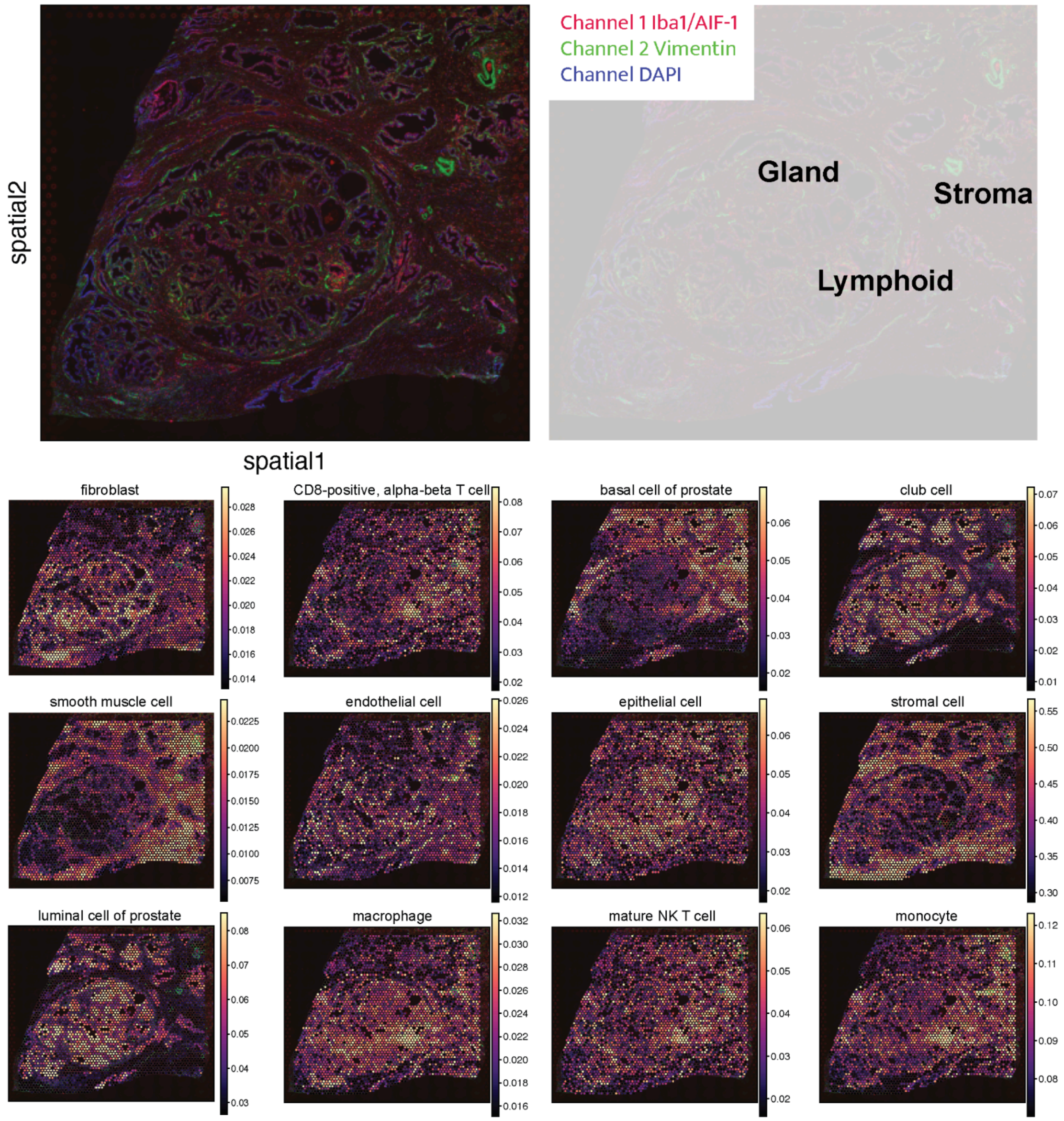


**Supplementary Figure 9: Deconvolution of Visium prostate cancer sample using Tabula sapiens reference model.** DestVI was used to deconvolve a human prostate cancer Visium sample. Top immunofluorescence image (see 10X data release) and legend in right plot. Tissue structures are annotated overlaid in the right plot. All other plots display the estimated cell-type proportions. There is a clear segregation between stroma (smooth muscle cells and stromal cells), lumen (luminal cells of prostate epithelial and club cells) and a lymphoid like structure in the center (CD8-positive, alpha-beta T cell, mature NK T cells, monocytes and macrophages).

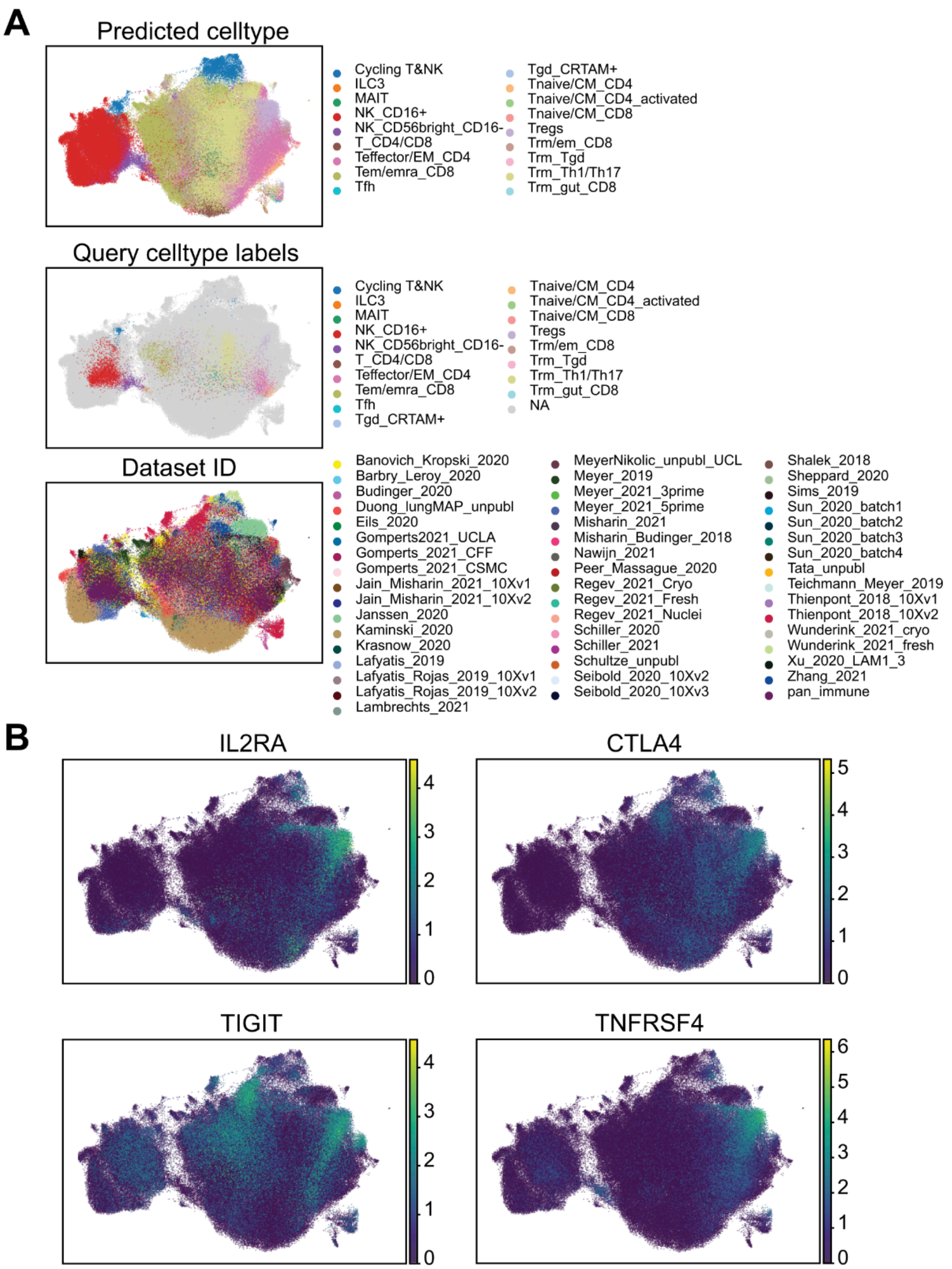


**Supplementary Figure 10: Label infusion from cross-organ immune cell atlas highlights TREG cells in HLCA. (A)** UMAP overlaid with additional covariates. First the predicted cell-types are displayed highlighting the split between NK and T cells. In the upper right cells are predicted to be TREGs. Query celltype labels are the input labels used for label transfer. Dataset ID is the source data ID. TREGs have contributions by a variety of source datasets, while Kaminski_2020 clusters very distinctly from other datasets. **b.** TREG markers from Figure 2 overlaid with UMAP from **(B)** Indeed TREG markers are positive in the region highlighted as TREG by label infusion. Both approaches identify similar cells.

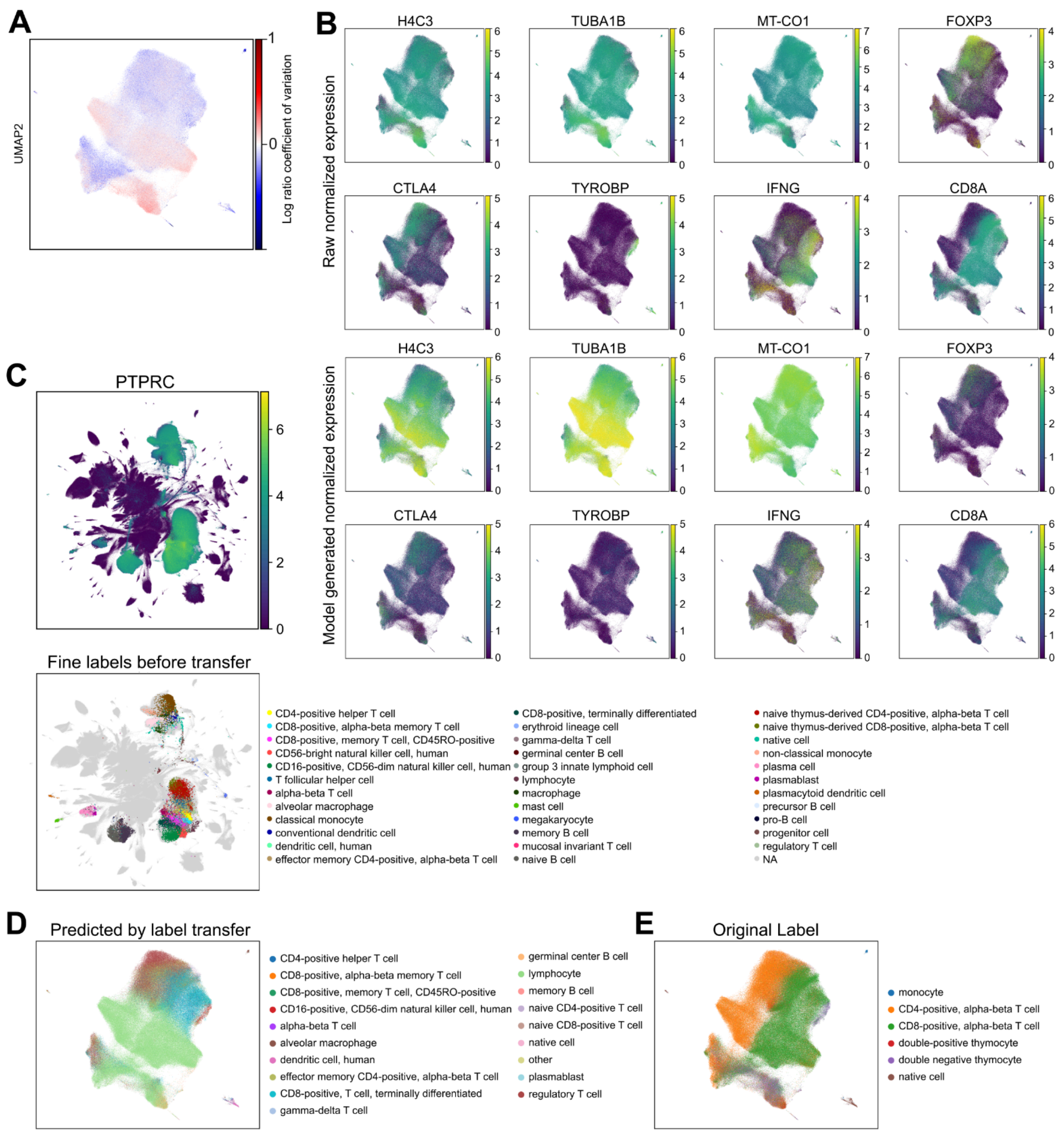


**Supplementary Figure 11: Query dataset analysis using CELLxGENE census model (A)** Cell-wise coefficient of variation overlaid on UMAP. Colored by the log-ratio of CV values between model generated and raw expression. Two clusters in the bottom with the largest discrepancy between model and raw. **(B)** Comparison of model-generated and raw expression values overlaid on UMAP. Display highlights more drastic differences than in **Figure 2** but model-generated expression and raw expression are expressed in similar regions. Displayed is expression after library size normalization and log1p transformation. **(C)** Top: PTPRC as a general immune cell marker. Displayed is raw expression after library size normalization and log1p transformation. Bottom: Labels of cross-organ immune cell atlas overlaid on UMAP (other datasets displayed in gray. Same embedding and coloring as in **Figure 4**, full legend is displayed here. **(D)** Cell-types are label transfer overlaid on UMAP, all cell-types predicted less than 100 times are summarized as *other*. **(E)** Original cell-type labels from the CAR T cells study.

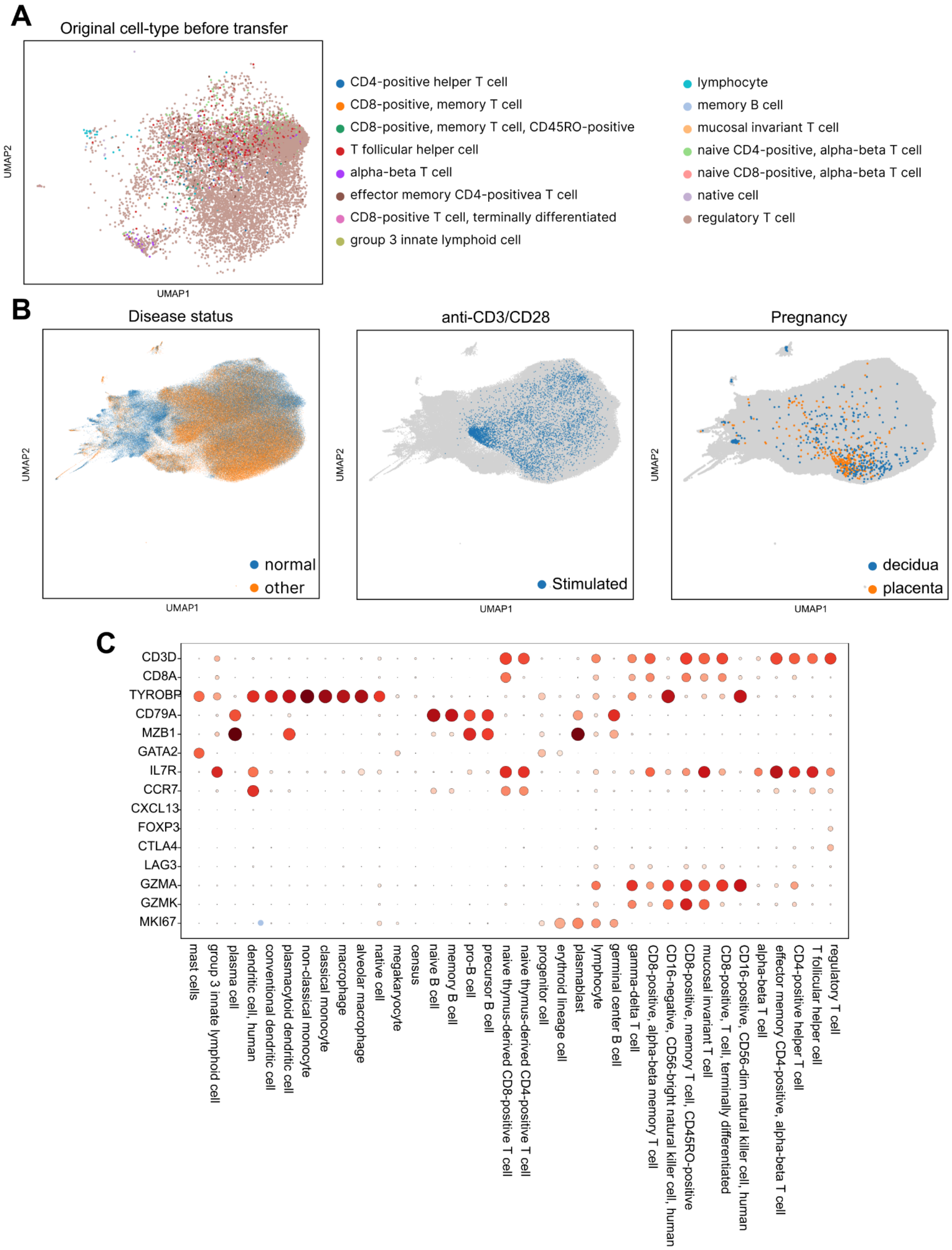


**Supplementary Figure 12: Annotating census dataset to generate novel insights. (A)** Cell-types of dataset used for label transfer overlaid on UMAP. In all regions cells are labeled in a majority *regulatory T cell*. **(B)** UMAP overlaid with disease status binarized to normal (healthy) and every other category. Enrichment of normal on the left side with naive TREGs. Highlighting a dataset annotated as normal but with *in vitro* stimulation leading to a high number of activated TREGs. Decidua and placenta are pregnancy associated tissues and contain a high number of activated TREGs. **(C)** Hand-selected marker genes for coarse cell-type groups highlighting that label transfer aligns with marker genes (CD3D for T cells, TYROBP for myeloid cells, CD79A and MZB1 for B cells**).**

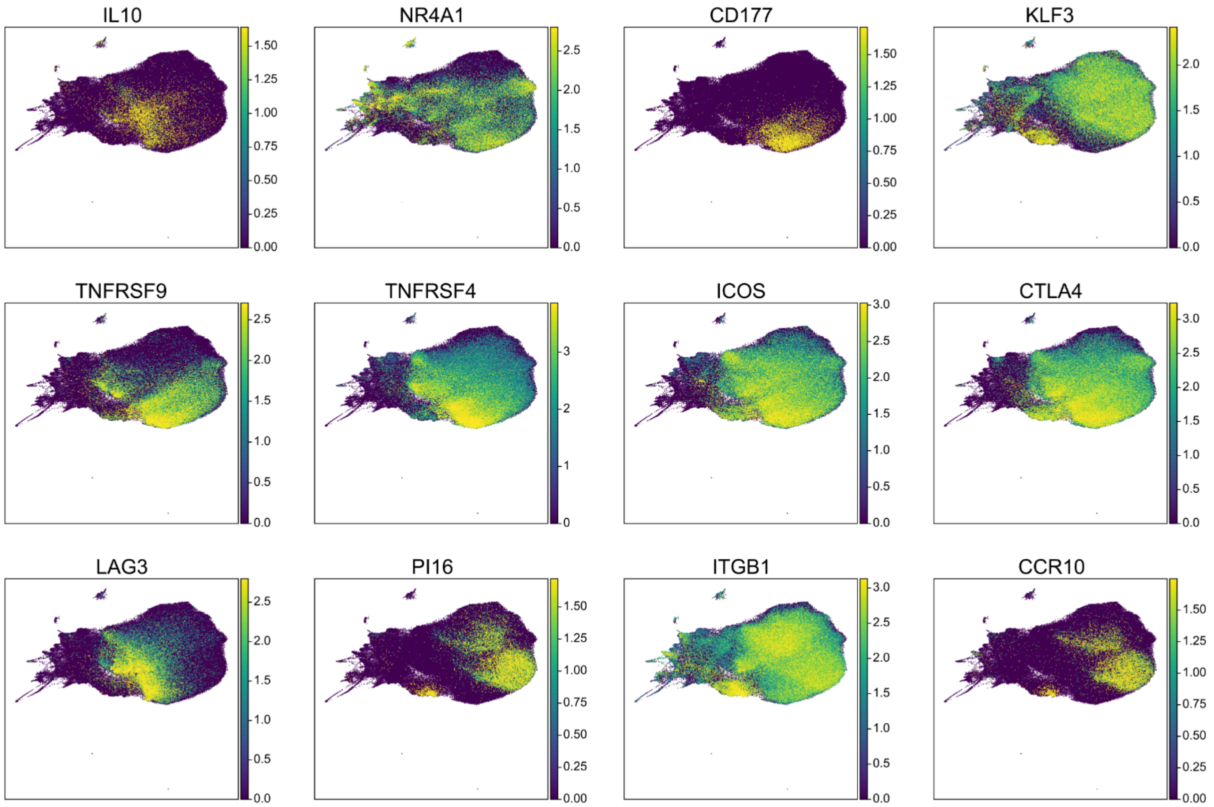


**Supplementary Figure 13: Heterogeneity of TREG cell states.** Activated TREGs are high in IL10, TNFRSF9, TNFRSF4, LAG3, CTLA4; similar states are observed in a subset of follicular helper T cells. PI16,ITGB1, CCR10 are higher expressed in memory Tregs. Displayed is the raw expression after library size normalization and log1p transformation.
